# Thymus formation in uncharted embryonic territories

**DOI:** 10.1101/2022.03.09.483697

**Authors:** Isabel Alcobia, Margarida Gama-Carvalho, Leonor Magalhães, Vitor Proa, Domingos Henrique, Hélia Neves

## Abstract

The thymus is a conserved organ among vertebrates, derived from the endoderm of distinct pharyngeal pouches (PP), whose location and number vary across species. Together with reports of sporadic ectopic thymus locations in mice and humans, this suggests that the potential to make a thymus resides in a broader region of the PP endoderm than previously ascribed.

Using the chick-quail chimera system, we explore this hypothesis and test the capacity of non-canonical pouches to participate in thymus formation. We further ask if the local mesenchyme of pharyngeal arches (PA) could also play a role in the regulation of thymus formation. After testing several embryonic tissue associations, we mapped the pharyngeal endoderm regions with thymus potential to the second and third/fourth pharyngeal pouches (2PP and 3/4PP). We further identified mesenchyme regions that regulate this potential to the 3/4 pharyngeal arches and to the dorsal region of the second arch, with positive and negative influences, respectively. Transcriptomic analysis of these tissues helped us revealing a common genetic program in the PP endoderm linked to thymus potential in addition to finding distinct signalling pathways involved in the cellular interactions with the mesenchyme of the pharyngeal arches that result in modulating this potential.

Together, these results provide new information about the initial specification of thymus primordia in the embryo that may contribute to improving the development of thymus organoid systems.

**Graphical abstract:** 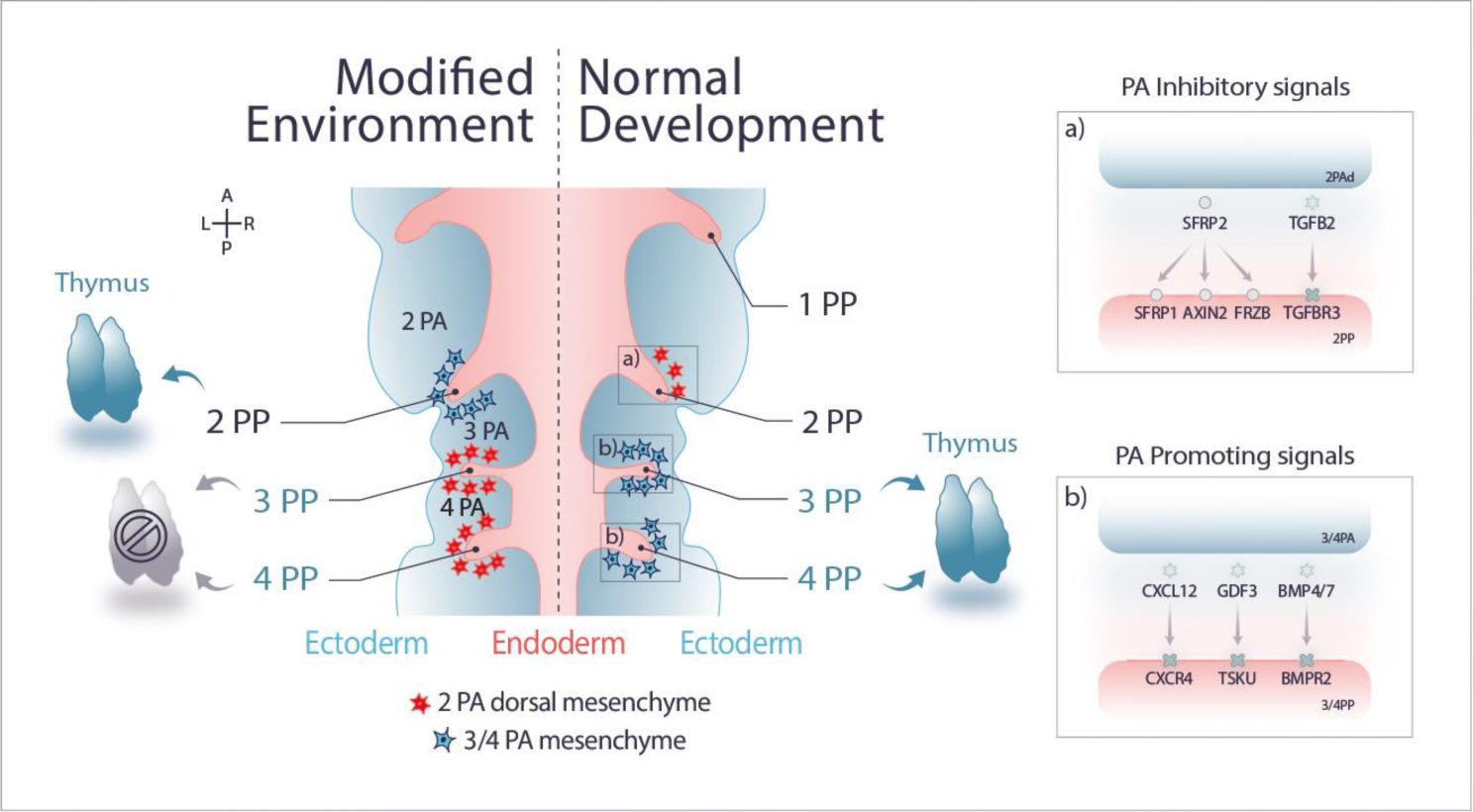

## Introduction

The thymus is a primary lymphoid organ essential for T-lymphocyte development and maturation. The generation of a self-tolerant and diverse repertoire of T-cell receptors will ensure an effective recognition and abolition of foreign antigens in the periphery without mounting a deleterious immune response to the body’s own tissues. This central immune tolerance is established by cellular interactions between differentiating lymphoid T-cells and a stromal microenvironment mainly formed by the thymic epithelium (TE) (Duah et al., 2021; Hamazaki, 2015).

The origin of TE rudiment in the endoderm of pharyngeal pouches (PP) was first demonstrated in 1975 using the chick-quail chimera system (Le Douarin and Jotereau, 1975). The thymus shares the same embryological origin with the parathyroid (PT) glands, an organ responsible for parathyroid hormone (Pth) production. The endoderm of the third PP (3PP) gives rise to the common T/PT primordium in mammals and birds, whereas the fourth PP (4PP) gives rise to the PT primordium only in humans and birds. In other vertebrates, such as frogs and reptiles, thymic rudiments are found in the second PP (2PP) and in the 2PP and 3PP (2/3PP), respectively [reviewed in (Rodewald, 2008)]. The final number and position of the thymus or thymi, are also variable among vertebrate species. While some mammals only have a thymus in the thoracic or cervical regions, others have thymi in both locations [reviewed in (Rodewald, 2008)]. In human and mice, the adult thymus is commonly positioned in the chest. Although rare, auxiliary cervical thymi can be found, as well as ectopic thymus inclusions in parathyroid glands (Dooley et al., 2006; Norris, 1938; Terszowski et al., 2006; Van Dyke, 1941).

Major efforts have been made to unveil the transcription factors and signalling molecules that affect thymus development prior to the specification of pharyngeal pouch endoderm into thymic fate. Though Fork-head Box N1 (*Foxn1*) transcription factor appears as the earliest TE marker (Nehls et al., 1994), it is dispensable for its specification, being only necessary for TE maintenance and differentiation (Blackburn et al., 1996; Nehls et al., 1996). Other involved transcription factors are *Tbx1*, *Pax1*, *Pax9*, *Hoxa3*, *Eya1*, and *Six1,* whose defects cause thymus aplasia or prevent thymus migration towards the chest [reviewed in (Figueiredo et al., 2020; Rodewald, 2008)]. Recent studies integrating single-cell transcriptomes and chromatin accessibility analysis provided new insight into the progression from embryonic endoderm cells into TE. Namely, *Grhl3* was found to be a putative new transcriptional regulator of the medullary TE sub-lineages (Magaletta et al., 2022; Park et al., 2020). In addition to the above-mentioned transcription factors, signalling pathways like bone morphogenetic protein (BMP), fibroblast growth factor (FGF), Wingless-int (Wnt), Notch, and Hedgehog were also shown to be involved in 3/4PP patterning and early phases of thymus organogenesis [reviewed in (Figueiredo et al., 2020)]. We have previously described the requirement for Notch signalling for TE specification and parathyroid epithelium differentiation in a manner that is Hedgehog dependent (Figueiredo et al., 2016). In addition, *Foxn1* expression was unexpectedly detected in the 2PP and its expression modulated by Hedgehog signalling, identical to what happens in the 3/4PP (Figueiredo et al., 2016). These findings suggest that *Foxn1*-expression domains in the endoderm of distinct pharyngeal pouches may embody a conserved potential to develop a thymus.

Given the known diversity of the PP location of thymic rudiments among vertebrates, we may envisage a potential conserved genetic program to "make a thymus" embodied in the endoderm of different PP, independent of their anatomical position. This program may be restricted by differential, yet unknown, molecular cues provided by the adjacent pharyngeal arch (PA) mesenchyme. To test this hypothesis, we took advantage of the modified chick-quail chimera system previously described by our lab (Figueiredo and Neves, 2018; Figueiredo and Neves, 2019; Figueiredo et al., 2016). We performed combinatorial heterospecific associations of PP endoderm with mesenchyme tissues from several embryonic locations and evaluated the potential to form a functional thymus. Additionally, transcriptome profiling by mRNA-sequencing (mRNA-seq) was performed in distinct endoderm and mesenchyme tissues of the embryonic pharyngeal arch region (PAR).

Our results show that non-canonical PP can generate a morphological and functional thymus upon interactions with permissive mesenchyme. This indicates the existence of a conserved program embodied in the endoderm of different PP, independently of their specific anatomical location. Furthermore, we identify the dorsal region of the 2PA mesenchyme as having inhibitory properties to thymus formation. Transcriptomic analysis revealed a shared genetic program between 2PP and 3/4PP endoderm, a previously unnoticed Hox-code in the prospective thymus rudiment as well as specific signalling pathway activities according to the distinct mesenchymal environments of PA.

## Results and discussion

### The endoderm of the 2PP has similar thymic potential to the 3/4PP endoderm

During chicken embryonic development, TE derives from the 3/4PP endoderm (canonical pouches). To evaluate the capacity of the endoderm of non-canonical pouches to form thymic epithelium, we started by removing them from their natural environment and associating them with a permissive mesenchyme, the somatopleura mesoderm (SM). The SM capacity to support thymus development at ectopic locations in the embryo was previously demonstrated when grafting 3/4PP endoderm into the body wall of a chicken embryo (Le Lièvre and Le Douarin, 1975; Neves et al., 2012). Using a modified chick-quail chimera system, 2PP endoderm isolated from quail (q) embryos at embryonic day 3 (E3) was *in vitro* associated with chicken (c) SM (E2,5), for 48h (Figueiredo and Neves, 2019; Neves et al., 2012). To evaluate the capacity of this heterospecific association to form a thymus, 48h-cultured tissues were then grafted into the chorioallantoic membrane (CAM) of a chicken embryo (E8) and allowed to develop *in ovo* for further 10 days (Figueiredo and Neves, 2018; Figueiredo et al., 2016) (Figure 1a). In parallel, associations of 3/4PP endoderm (qE3) with SM (cE2,5) were used as positive control (Neves et al., 2012). Morphological analysis of explants was performed by conventional histology and immunohistochemistry, as previously described (Figueiredo and Neves, 2018; Figueiredo et al., 2016). Endoderm-derived cells were identified by immunohistochemistry using a quail-specific antibody, mAb Quail PeriNuclear (QCPN).

**FIGURE 1.**
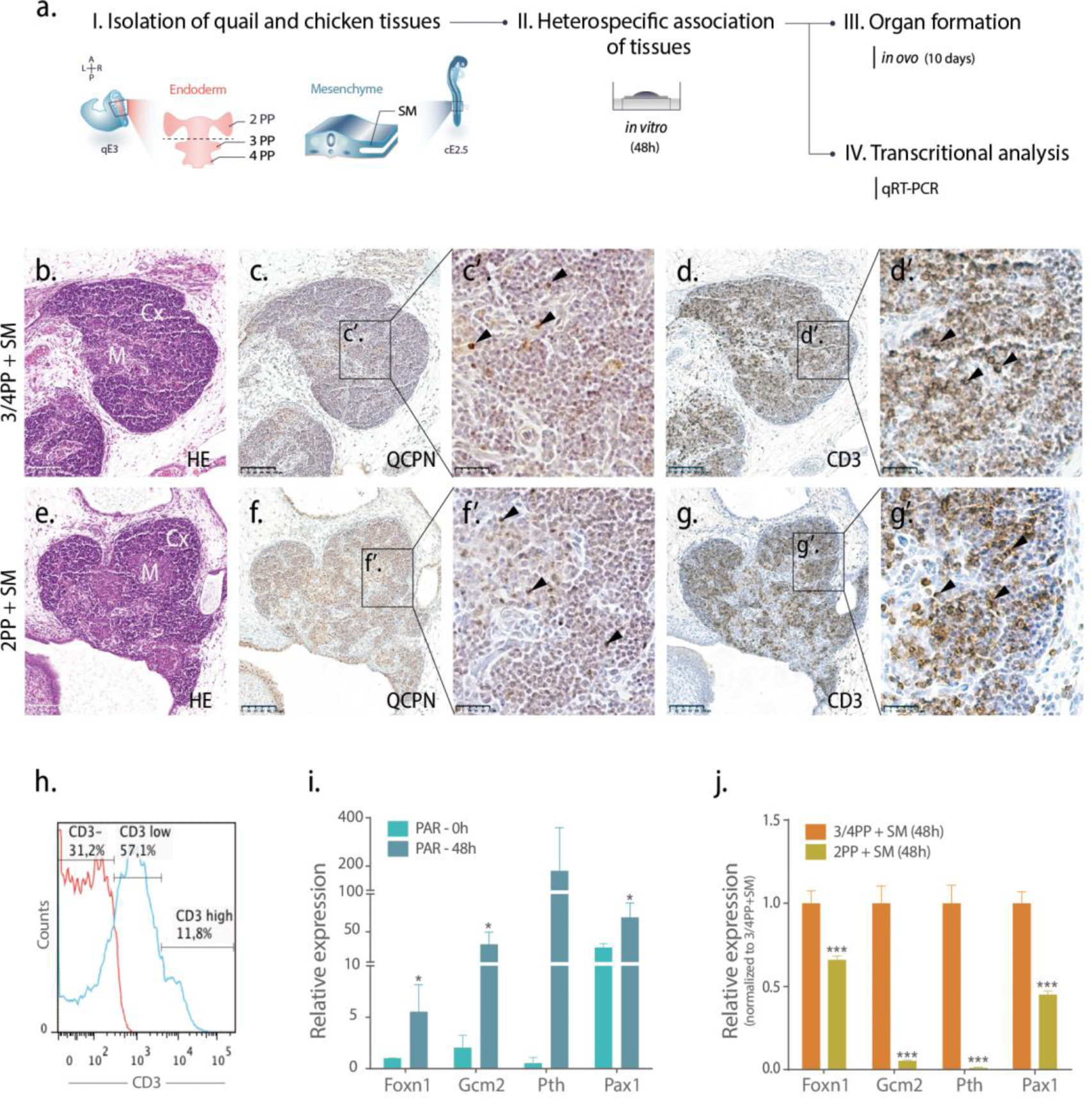
| Thymus formation in PP endoderm-derived explants. a) Schematic representation of *in vitro* and *in ovo* experiments wherein quail 2PP or 3/4PP endoderm was associated with chicken SM and cultured for 48h. Cultured tissues were then either grafted and grown *in ovo* for an additional 10 days onto the CAM at cE8 or were analysed by qRT-PCR. b-g) Serial sections of thymi obtained in PP endoderm-derived explants stained with H&E (b and e), and immunodetected with QCPN (quail cells) (c and f) and CD3 (T-cells) (d and g) antibodies. Magnified images of c, d, f and g in c’, d’, f’ and g’, respectively. Arrowheads point immunostaining positive cells for QCPN (quail TE) and for CD3 (chicken T-cells differentiated in the thymus). h) Histogram of surface CD3 expression on thymocytes isolated from chicken thymus at E17. Unstained and CD3-stained thymocyte are depicted in red and in blue, respectively. i-j) Gene-expression levels of *Foxn1*, *Gcm2*, *Pth* and *Pax1* by qRT-PCR. The expression of each transcript was measured as a ratio against Hprt transcript expression level and expressed in arbitrary units. qRT-PCR analysis was performed in PAR explants [freshly isolated (PAR-0 h) and cultured (PAR-48 h)] (i) and in 48h-cultured tissues (j). In (j), the expression levels of transcripts were normalized to the control condition, 3/4PP endoderm and SM cultured for 48h (each transcript in control=1). A, Anterior; CAM, chorioallantoic membrane; cE, chicken embryonic day; L, Left; P, Posterior; Cx, cortex; M, medulla; PAR, Pharyngeal Arch Region; PP, Pharyngeal Pouch; qE, quail embryonic day; SM, somatopleura mesoderm; R, right. Scale bars, 100μm.

Thymi were detected in all explants examined, derived from both endoderm of 3/4PP (n=5/5) and 2PP (n=6/6)(Table 1). The 2PP endoderm-derived explants contained chimeric thymi with cortical and medullary compartments (Figure 1e), as in control explants (Figure 1b)(Neves et al., 2012). Quail-derived TE displayed a reticular architecture (QCPN+, Figure 1c and Figure 1f) and was colonised by lymphoid progenitor cells of donor origin (chicken). To explore T-lymphoid differentiation, several T-cell markers were analysed by flow cytometry (Figure S1). Due to antibody cross-reactivity limitations, this study was performed in isolated thymocytes from chicken embryos. CD3 expression was observed from day 11 onwards, reaching around 70% of T-cell population by day 17 of development (Figure 1h). As in normal development, chimeric thymic showed a majority of CD3+ thymocytes in both conditions (Figures 1d and 1g). Together, these results suggest a similar potential of non-canonical 2PP and canonical 3/4PP endoderm to specify TE.

**Table 1.** Formation of chimeric thymus and parathyroids in CAM-explants.

| **Heterospecific associations of tissues** | **Thymus** | **Parathyroids** |
| --- | --- | --- |
| 3/4PP endoderm + Somatopleura mesoderm | 100% (n=5/5) | 80% (n=4/5) |
| 2PP endoderm + Somatopleura mesoderm | 100% (n=6/6) | 67% (n=4/6) |
| 2PP endoderm + 3/4 PA mesenchyme | 100% (n=3/3) | 67% (n=2/3) |
| 3/4PP endoderm + 3/4 PA mesenchyme | 100 % (n=3/3) | 100 % (n=3/3) |
| 3/4PP endoderm + 2 PA mesenchyme | 100% (n=3/3) | 67% (n=2/3) |
| 3/4PP endoderm + Ventral territory of 2 PA mesenchyme | 100 % (n=3/3) | 67% (n=2/3) |
| 3/4PP endoderm + Dorsal territory of 2 PA mesenchyme | 20% (n=1/5) | 100% (n=5/5) |

n= number of samples with organs per total number of samples.

It is worth mentioning that no thymus was formed when 1PP endoderm was associated with SM (n=6/6). Only small clusters of epithelial cells with occasional CD3+ cells were observed in the explants, indicating a residual potential of this pouch endoderm to attract lymphoid cells (Figure S2a-e).

Given the common primordium origin of the thymus and PT glands epithelia, we further analysed the presence of parathyroid tissue in endoderm-derived explants. PT glands were formed in explants derived from 2PP endoderm (not shown), although with lower efficiency (n=4/6) than in 3/4PP (n=6/6)(Table 1)(Neves et al., 2012). Together, the data suggest a conserved mechanism for TE and PT formation within distinct pouches.

To further assess 2PP endoderm potential, heterospecific associations of tissues were cultured for 48h and analysed for the expression levels of TE- and PT-related markers, *Foxn1* (Neves et al., 2012) and *Gcm2/Pth* (Grevellec et al., 2011; Neves et al., 2012), respectively. Expression analysis was extended to *Pax1*, encoding a transcription factor known to be involved in the morphogenesis of the pouches and thymus and PT formation (Dietrich and Gruss, 1995; Guo et al., 2011; Su et al., 2001; Wallin et al., 1996).

We started by *in vitro* monitoring the gene expression levels in the developing pharyngeal arch region (PAR) without disturbing pharyngeal pouches environment. For that, the ventral portion of PAR from 2PA to 4PA at qE3, was dissected and cultured *in vitro* for 48h (Figueiredo et al., 2016). Predictably, very low levels of expression for TE- and PT-related markers were detected in freshly dissected tissues (PAR-0h) (Figure 1i) (Figueiredo et al., 2016). However, 48h-cultured tissues exhibited a strong and significant increase in expression levels of the four genes when compared to fresh tissues (Figure 1i) (Figueiredo et al., 2016). Then, we explored the expression levels of *Foxn1, Gcm2/Pth* and *Pax1* in PP endoderm isolated and cultured with SM for 48h. Transcript levels of the four genes were quantified in cultured tissues and normalized to the control condition (each transcript=1), the association of 3/4PP endoderm with SM. We observed a significant lower expression of the four genes in the 2PP condition (Figure 1j), revealing a distinct ability of each pouch endoderm to express TE and PT-related genes. The 40% and 60% decreases in *Foxn1* and *Pax1 in vitro* expression (Figure 1j) point to a developmental delay of 2PP endoderm in acquiring thymic markers. However, given that normal thymus formation occurs in 2PP-derived explants with comparable efficiency to controls (Figure 1e-g and Table 1), this delay must be overcome during long-term development. The almost absence of PT-specific gene expression in 2PP endoderm after 48h of culture (5% *Gcm2* and 1% *Pth*), combined with reduced efficiency in PT formation (Table 1), suggests a weaker potential of 2PP endoderm to form PT when compared to canonical 3/4PP endoderm.

Taken together, these data point to the existence of a conserved program to form a thymus embodied in the endoderm of canonical 3/4PP and non-canonical 2PP.

### The 2PP and 3PP endoderm share a conserved genetic program

Considering the capacity of 2PP endoderm to give rise to TE, we then asked if these endoderm tissues could share a common genetic program with the 3/4PP endoderm. To evaluate this, the study progressed by employing a mRNA-Seq approach to obtain the transcriptome profiling of the pouches. Briefly, the anterior endoderm region at qE3 was isolated (Figueiredo and Neves, 2019; Neves et al., 2012) and separated in three regions: 2PP, 3/4PP and the central territory of anterior endoderm (pharynx), the latter used as a negative control to pouches potential (Figure 2a). A pool of each type of tissue isolated from 15 to 20 embryos was used to generate a biological sample. Total RNA was isolated from three biological replicate samples and used to generate libraries for Illumina paired-end mRNA-seq. High quality sequencing datasets were obtained for all samples, with an average 7 million reads mapping uniquely to the quail genome, corresponding to the detection of ∼14 500 genes (Supplementary Dataset 1). The generated tree of differentially expressed (DE) genes heatmap showed a closer similarity between the endoderm of the pouches, than with the pharynx (the complete list of DE genes can be found in Supplementary Dataset 2). Association of genes based on their expression trends revealed six clusters (Figure 2b). Cluster 1 displayed an enrichment of highly and specifically expressed transcripts in 3/4PP endoderm. Of those, *Gcm2*, *Eya2*, *Pax1*/*9*, *Hoxa3*, *Mafb*, *Gata3*, are transcription factors known to be involved in 3/4PP patterning and common T/PT primordium specification in various animal models [reviewed in (Figueiredo et al., 2020)]. Additionally, *Jag1*, a Notch ligand, was also shown to be more expressed in the 3/4PP endoderm (Figueiredo et al., 2016). Clusters 2 and 5 show transcripts more expressed in the 2PP endoderm while clusters 4 and 6 presented a pool of transcripts that are more expressed in the pharynx. Interestingly, clusters 2 and 5 were enriched in transcripts of Wingless-int (Wnt) pathway ligands suggesting a possible autocrine or paracrine signalling modulation of 2PA environment by 2PP endoderm cells. GO-term enrichment analysis was performed on each cluster to gain insight into the function of each class of DE genes (Figure 2c and Supplementary Dataset 3). We found that genes from 2PP and pharynx endoderm clusters (Clusters 2-5 and 4-6, respectively) were involved in a few Biological Process (BP) categories. In contrast, an enrichment in several GO-BP categories was found in the “3/4PP endoderm cluster” (Cluster 1) such as Positive regulation of cell migration, Cell population proliferation, Morphogenesis of an epithelium, Tube morphogenesis, and other Go-terms related to signalling pathways. In particular, the Go-term FGFR signalling pathway was linked to cluster 1. Within this cluster, we found FGF ligands such as Fgf3, Fgf18, and Fgf19. The expression of these Fgfs was validated by whole mount *in situ* hybridization in chicken embryos at the same developmental stage (Figure 2d). While transcripts were detected in all PP endoderm, the stronger hybridization signals were always observed in the 3/4PP endoderm, the most recently formed pouches. This suggests a shared genetic program that is chronologically expressed during pouches morphogenesis. Accordingly, Cluster 3 represents a pool of transcripts commonly expressed in the endoderm of 2PP and 3/4PP, enriched in GO-BP categories such as Pattern specification process and Pharyngeal system development, further supporting the concept that pouches share common basic developmental processes.

**FIGURE 2.**
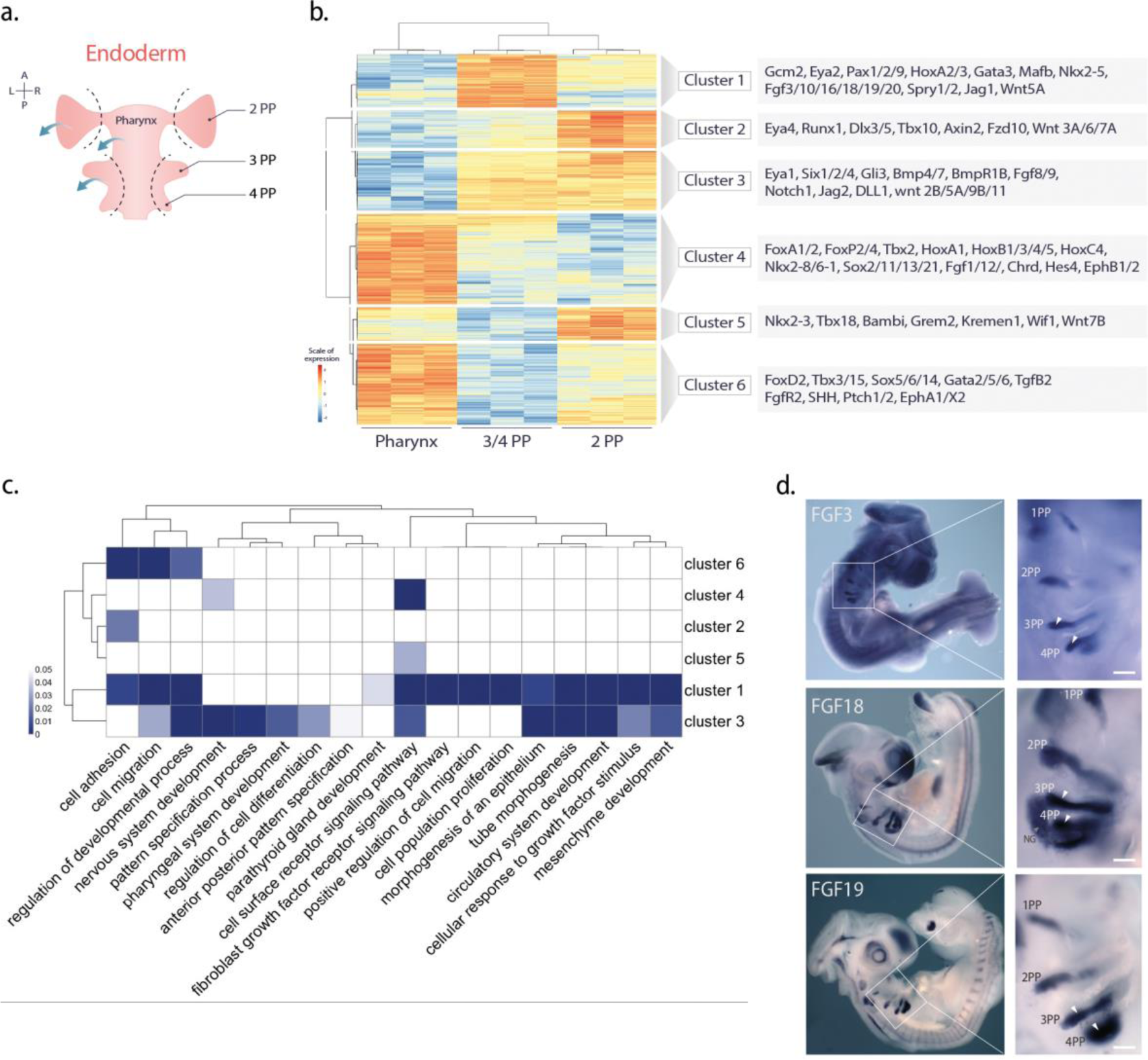
| Transcriptional profiles of pharyngeal endoderm tissues. a) Schematic representation of distinct endoderm tissues isolated from the embryonic pharyngeal region at qE3. b) Unsupervised clustering of differentially expressed genes in the pharynx, 3/4PP and 2PP endoderm (fold change >0.6 and <−0.6, and *P*-value <0.01), highlighting the grouping of genes into six different clusters of expression. Known markers belonging to each cluster are shown in the boxes on the right. The expression values reported in the heatmap correspond to row-scaled (Z-score), log transformed count data (see colour scale on the left of the plot). c) Heatmap showing the results of the Biological Process GO-term enrichment analysis performed on the six gene clusters shown in b. The heatmap reports only a set of the significantly enriched categories (FWER <0.05 in at least one cluster), selected in order to reduce redundancy. The colour intensity in each cell represents FWER according to the scalebar on the left. d) Expression of *FGF3*, *FGF18 and FGF19* detected by *in situ* hybridization in the chicken pharyngeal region at embryonic day 3.5. White arrowheads point to high levels of expression in pharyngeal pouches. Magnified images of a-c in a’-c’, respectively. Scale bars, 100 μm. A, Anterior; L, Left; P, Posterior; PP, Pharyngeal Pouch; R, right.

### Genetic signature of pouch endoderm with thymic potential

To obtain some insights regarding the genetic programs that regulate the thymic potential of different pouches, we deepen our analysis of the distinct transcriptome profiles of endodermal tissues. We identified 1224 and 1085 differentially expressed (DE) genes in the 2PP or 3/4PP endoderm as compared with the pharynx, respectively (the complete list of DE genes can be found in Supplementary Dataset 2). Of these, we found 555 and 492 upregulated transcripts (log2FC >0.6, adj pval <0.01) in the 2PP and 3/4PP endoderm when compared to the pharynx, respectively. Additionally, these two groups had in common 262 upregulated transcripts (Figure 3a). Then, we deepened our analysis to identify players of genetic programs that may be specifying thymic competence in endoderm progenitors of the pouches. Upregulated transcription factors and regulators (TFs&Rs) within this set of genes were identified by manual examination of gene annotations using Ensembl genome and RefSeqGene browsers (Figure 3a).

**FIGURE 3.**
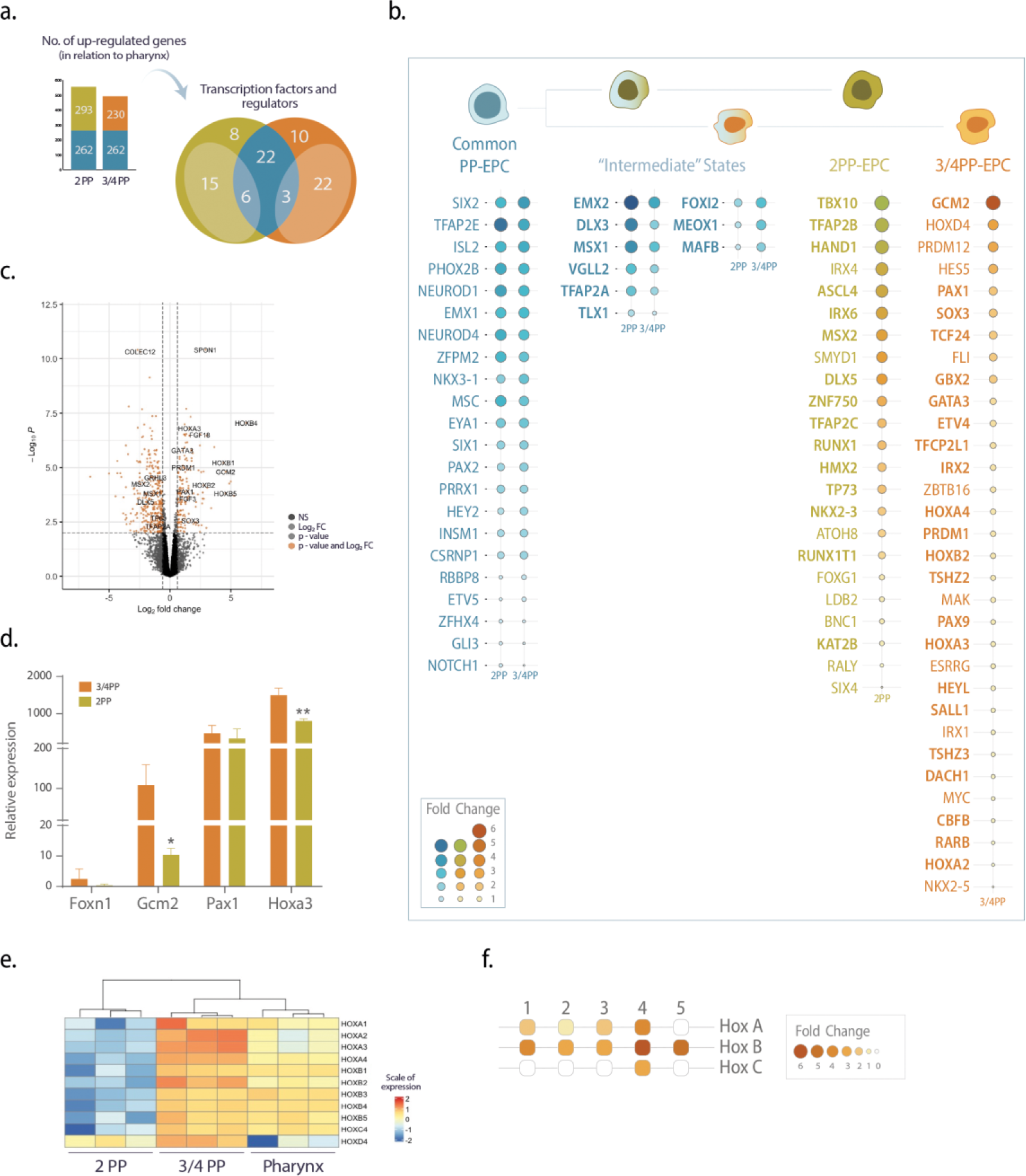
| Differential gene expression analyses reveal discrete transcriptomic signature of 2PP and 3/4PP endoderm. a) Bar plot depicting the number of upregulated genes in 2PP and 3/4PP transcritomes in relation to pharynx transcriptome and Venn diagram depicting the overlap of a selection of transcription factors and regulators. b) Schematic representation of the genetic profile of distinct PP progenitors considering the upregulated transcription factors and regulators founding in PPs. The dot size and color intensity represent fold change in expression level. Different colors correspond to putative cell states represented on the top of the panel. Bold highlighted genes identify upregulated transcripts in relation to the other PP as well as to the pharynx. c) Volcano plot showing the results of the differentially expressed analysis between 3/4PP and 2PP endoderm. Orange dots highlight transcripts passing the log2FC and adj pvalue cut-offs (adj pval <0.01 and |log2FC|>0.6); grey dots highlight transcripts passing only one of the cut-offs. d) Gene-expression levels of *Foxn1*, *Gcm2*, *Pax1* and *Hoxa3* by qRT-PCR. The expression of each transcript was measured as a ratio against *Hprt* transcript expression level and expressed in arbitrary units. qRT-PCR analysis was performed in freshly isolated 2PP and 3/4PP endoderm. e) Heatmap showing differentially expressed Hox family genes in the pharynx, 3/4PP and 2PP endoderm (|log2FC|>0.6, and adj pval <0.01). The expression values reported in the heatmap correspond to row-scaled (Z-score), log transformed count data (see colour scale on the right of the plot). f) Drawing depicting upregulated transcripts of Hox-gene family members (A-D and 1-5) in 3/4 PP endoderm when compared to the 2PP endoderm (see colour scale on the right of the plot). NS, Non-Significant; PP, Pharyngeal Pouch.

Given that the developmental potential of 2PP endoderm mimics that of 3/4PP, it is foreseen that the pool of commonly upregulated TFs&Rs may represent a genetic program for a putative common PP endoderm progenitor (Figure 3b). We found 22 transcripts encoding TFs&Rs that show no DE between pouches (Figure 3b). Of these, *Eya1/Six1* (Li et al., 2003), *Notch1*/*Gli3* (Figueiredo et al., 2016), and *NeuroD1* (Bautista et al., 2021; Park et al., 2020), have been shown to play critical roles in the early phases of T/PT organogenesis in mouse, chicken, and human (Figure 3h). Despite being implicated in several developmental processes in different species, the remaining 17 (out of the 22) TFs&Rs have no known role in thymus formation. Only *Insm1* was found to be expressed in the human medullary TE, particularly around Hassall’s corpuscles (Kashima et al., 2022) and mutant Etv5 mice display decreased adult thymus weight (Gutierrez-Aguilar et al., 2015).

Some of the commonly upregulated TFs&Rs were also DE between pouches, suggesting a genetic signature of "intermediate” stages of commitment into the distinct endoderm pouch progenitors (Figures 3a and 3b). Specifically, 3 transcripts encoding the TFs&Rs, *Meox1*, *Mafb,* and *Foxi2*, were upregulated in 3/4PP endoderm when compared to 2PP endoderm, signing a putative intermediate stage of commitment into 3/4PP endoderm progenitors (Figure 3b). *Meox1* and *Mafb* were mainly identified in parathyroid development (Kamitani-Kawamoto et al., 2011; Magaletta et al., 2022), while *Foxi2* was shown to be expressed in PA mesenchyme of chicken embryos (Khatri and Groves, 2013). In contrast, 6 transcripts encoding TFs&Rs were found to have higher expression in 2PP endoderm (*Dlx3*, *Emx2*, *Msx1*, *Tfap2A*, *Vgll,* and *Tlx1*), which may hint to a comparable "intermediate" stage of commitment into 2PP endoderm progenitor (Figure 3b). It’s interesting to note that Tfap2A functions upstream of Msx1 and Mafb in neural crest progenitors, (Enkhmandakh and Bayarsaihan, 2015), and mutations in this gene result in branchio-oculo-facial syndrome.

Lastly, our analysis revealed a set of 23 and 32 transcripts encoding TFs&Rs specifically upregulated in 2PP or 3/4PP endoderm as compared with the pharynx, respectively (Figures 3a and 3b). These two gene sets might be viewed as a further step into the gene cascade that regulates the acquisition of 2PP and 3/4PP identities. To refine these identities, we further compared the transcriptomes of distinct pouches’ endoderm (Figure 3c) and we identified 518 DE transcripts between the 2PP and 3/4PP endoderm (the complete list of DE genes can be found in Supplementary Dataset 2). Of these, 165 and 353 transcripts were upregulated in 3/4PP and 2PP endoderm, respectively. The extracellular matrix protein, *spondin 1* (*Spon1*), was the highest expressed transcript in the pool of up-regulated genes in 3/4PP endoderm (Figure 3c), suggesting a previously unrecognised role of this extracellular matrix protein in thymus formation. This result is concordant with our pilot microarray dataset (repository Zenodo) and other studies (GEISHA ISH Analysis). From the total of 34 upregulated transcripts encoding TFs&Rs, 22 were identified in the upregulated set of genes when compared to the pharynx (Figures 3a and b). Worth mentioning the transcripts encoding TFs&Rs, *Gcm2*, *Hoxa3*, *Gata3*, *Pax1,* and *Pax9*, above identified in cluster 1 (Figure 2b), are known to be involved in 3/4PP patterning and common T/PT primordium specification in various animal models [reviewed in (Figueiredo et al., 2020)]. Other transcripts such as *Hoxa2*, *Gbx2*, *Sall1,* and *Prdm1* were recently found in immature pharyngeal cells and thymic epithelial progenitors while *Hoxb2* was associated with ultimobranchial body formation in 4PP endoderm (Magaletta et al., 2022). Of the remaining TFs&Rs, *Sox3*, which is known to be necessary for PP formation and subsequently craniofacial morphogenesis (Rizzoti and Lovell-Badge, 2007), may play a yet-unknown role in the latter stages of posterior pouches maturation.

From the remaining pool of 10 transcripts encoding TFs&Rs exclusively upregulated in 3/4PP in comparison to the pharynx, but not to the 2PP endoderm, it is of note that *Nkx2*.*5* and *Myc* were found in immature pharyngeal cells and thymic epithelial progenitors (Magaletta et al., 2022). The Notch-target transcript *Hes5* was also found in this set of genes, suggesting the involvement of Notch activity in these early stages of thymus organogenesis. Indeed, Notch signalling was shown to be required for TE specification and PT epithelium differentiation and, *Hes5* expression was detected in the developing 3/4PP endoderm (Figueiredo et al., 2016).

The expression of *Hoxa3*, *Gcm2,* and *Pax1* obtained by RNAseq data was validated by qRT-PCR (Figure 3d) in freshly isolated endoderm of 2PP and 3/4PP at qE3. Transcript levels for *Gcm2* and *Hoxa3* were considerably higher in 3/4PP endoderm than 2PP endoderm, as observed in RNAseq data. *Pax1* transcripts levels were slightly higher in 3/4PP endoderm, with a similar tendency to RNAseq data. *Foxn1* transcript levels were barely detected, as previously described (Neves et al., 2012), and in agreement with no transcript detection of *Foxn1* by RNAseq.

The analysis of DE genes between transcriptomes of distinct pouches’ endoderm progressed to the evaluation of 2PP endoderm up-regulated transcripts (Figure 3c). From the total of 40 upregulated transcripts encoding TFs&Rs, 15 were identified in the upregulated set of genes when compared to the pharynx (Figures 3a and b). In this pool of TFs&Rs, *Grhl3*, *Tp63*, *Tfap2A*, *Msx1*, *Msx2*, *Tbx10*, *Dlx5,* and *Sox9* are known to be involved in orofacial cleft disease (Reynolds et al., 2019; Reynolds et al., 2020).

To finish our analysis of the distinct endoderm transcriptomic profiles, we have noticed DE genes belonging to TFs Hox-family of genes, known to define position identity in the developing embryo (Young and Deschamps, 2009). The heatmap tree generated by the 11 DE transcripts encoding Hox-genes, Hoxa1-4, Hoxb1-5, Hoxc4 and Hoxd4, showed a closer similarity between the 3/4PP endoderm and pharynx, than with the 2PP endoderm (Figure 3e). A set of 10 Hox genes were upregulated in 3/4PP endoderm when compared to 2PP endoderm (Figures 3e and 3f). *Hoxa3* is involved in 3/4PP patterning and common T/PT primordium specification [reviewed in (Figueiredo et al., 2020)] while *Hoxb1*, *Hoxb2*, *Hoxb4*, and *Hoxb5* were recently associated with ultimobranchial body formation in 4PP endoderm (Magaletta et al., 2022). The remaining 5 Hox genes have not yet been attributed functions related to thymus formation. The upregulated Hox genes in 3/4PP (Figure 3f) points to a discrete boundary between the pouches, which may be essential for anterior-posterior (A-P) pouch identity and proper specification of the thymus.

### The mesenchyme of PA modulates the thymic potential of PP endoderm

The ability of 2PP endoderm to develop into a thymus in the presence of ectopic mesenchyme demonstrated not only the presence of thymic potential in this region but also the importance of signals from local PA mesenchyme for regulating such potential in PP endoderm. The typical absence of thymus in 2PP may indicate that the underlying 2PA mesenchyme does not support thymus development or may even offer a local environment that restricts its formation. The 3/4PA mesenchyme, on the other hand, is anticipated to produce signals that promote thymus development in the underlying endoderm.

To start testing these hypotheses, 2PP endoderm was first isolated from chicken embryos and associated *in vitro* with 3/4 PA mesenchyme, followed by growth *in ovo* (Figure 4a), as described above. As anticipated, we observed thymi in the 2PP endoderm-derived explants having normal morphology with most thymocytes expressing the CD3 marker and presenting a TE of quail origin (Figures 4b-d). The efficiency of thymus formation was identical to control explants (3/4PP endoderm-derived) (Table 1 and Figures 4e-g).

**FIGURE 4.**
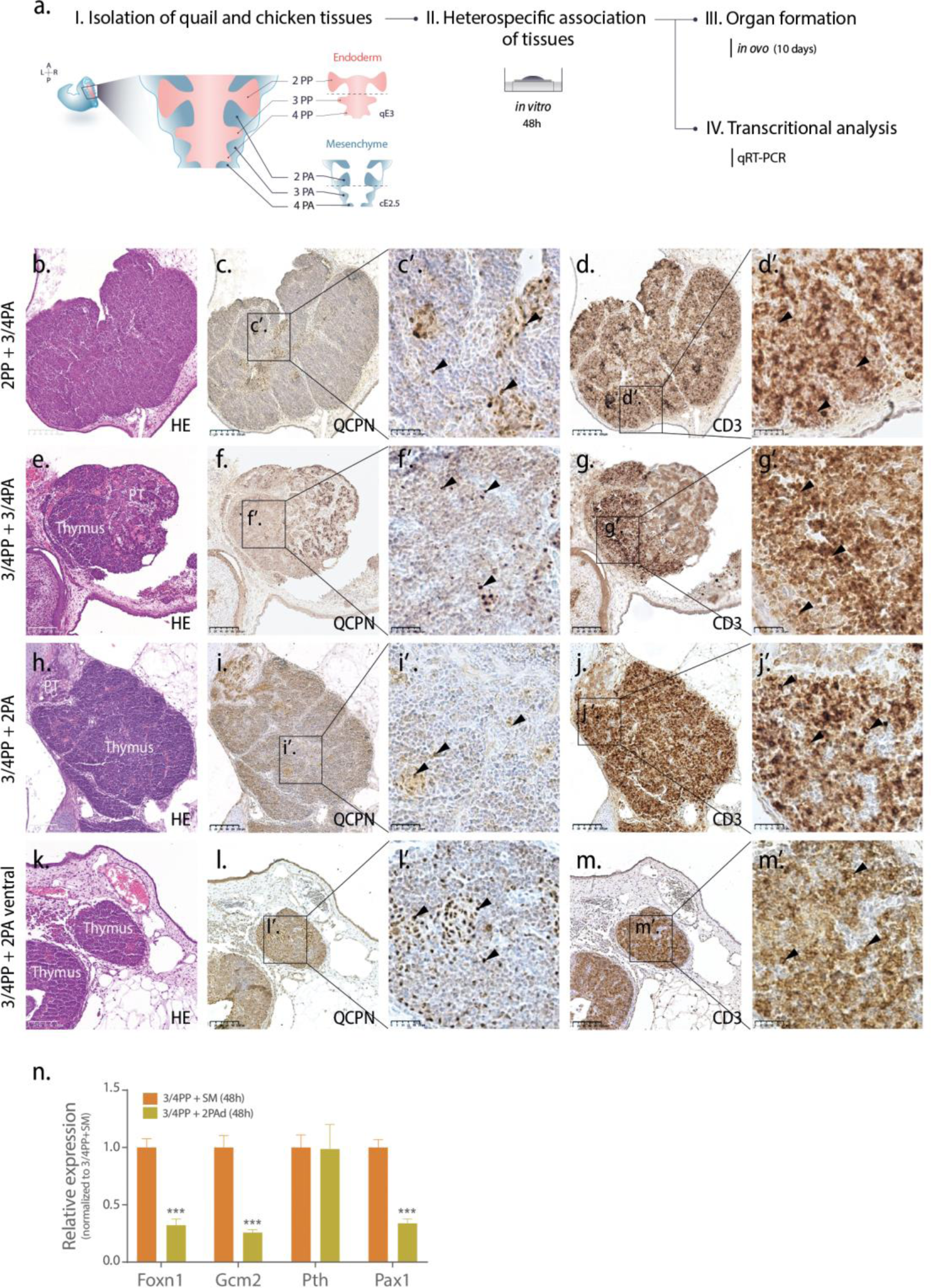
| Thymus formation from association of PP endoderm and PA mesenchyme. a) Schematic representation of *in vitro* and *in ovo* experiments wherein quail 2PP or 3/4PP endoderm was associated with chicken 3/4PA or 2Pad mesenchyme and cultured for 48h. Cultured tissues were then either grafted and grown *in ovo* for an additional 10 days onto the CAM at cE8 or were analysed by qRT-PCR. b-k) Serial sections of thymi obtained in PP endoderm-derived explants stained with H&E (b, e, h and k), and immunodetected with QCPN (quail cells) (f, f, i and l) and CD3 (T-cells) (d, g, j and m) antibodies. Magnified images of c, d, f, g, i, j, l and m in c’, d’, f’, g’, I’, j’, l’ and m’ respectively. Arrowheads point immunostaining positive cells for QCPN (quail TE) and for CD3 (chicken T-cells differentiated in the thymus). n) Gene-expression levels of *Foxn1*, *Gcm2*, *Pth* and *Pax1* by qRT-PCR. The expression of each transcript was measured as a ratio against Hprt transcript expression level and expressed in arbitrary units. qRT-PCR analysis was performed in 48h-cultured tissue associations of 3/4PP endoderm and 2PAd mesenchyme. The expression levels of transcripts were normalized to the control condition, co-culture of 3/4PP endoderm and SM for 48h (each transcript in control=1). A, Anterior; CAM, chorioallantoic membrane; cE, chicken embryonic day; L, Left; P, Posterior; PA, Pharyngeal Arch; PAd, Dorsal region of Pharyngeal Arch; PP, Pharyngeal Pouch; qE, quail embryonic day; SM, somatopleura mesoderm; R, right. Scale bars, 100μm.

To test for the presence of thymus-repressive signals in 2PA mesenchyme, this tissue was associated with 3/4PP endoderm (Figure 4a). Unexpectedly, normal thymi were formed in all explants derived from this type of association (Table 1 and Figures 4h-j). Considering that thymic rudiments in chicken embryos emerge at the dorsal tip of 3/4PP, we then asked whether the mesenchyme of the arches could have regionalized properties along the dorsoventral axis. To explore this hypothesis, the 2PA mesenchyme was isolated and then sectioned into dorsal (2PAd) and ventral (2PAv) regions, by microsurgical procedures (Figueiredo and Neves, 2019). These regions were associated with 3/4PP endoderm. Explants derived from 2PAv mesenchyme and 3/4PP endoderm associations displayed normal thymus formation (n=3/3, Figures 4k-m). In contrast, only one in six CAM-explants formed thymi when 3/4PP endoderm was associated with 2PAd mesenchyme, revealing the inhibitory properties of the latter to thymus formation (Table 1). Of notice, 1PA mesenchyme also showed permissive properties to thymus formation: tissue associations of 3/4PP endoderm with 1PA mesenchyme were able to form thymi (n=5/6, Figure S1).

To pursue the evaluation of 2PAd mesenchyme properties, heterospecific associations with 3/4PP endoderm were cultured for 48h and examined by qRT-PCR (Figure 4a), as detailed above. The 3/4PP endoderm cultured for 48h with SM was used as a control condition (each transcript = 1). A significant reduction of *Foxn1* transcript levels (>80%) was observed in 3/4PP endoderm grown with 2PAd mesenchyme when compared to control (Figure 4n). The results point to an early and perhaps irreversible inhibition of *Foxn1* expression when the 3/4PP endoderm is placed in contact with 2PAd mesenchyme. As an outcome, an important inability to form a thymus was observed in this condition (Table 1). Though a similar repression was observed for *Gcm2* expression, no changes in *Pth* transcript levels were detected when 3/4PP endoderm was grown with either 2PAd or SM (Figure 4n). Explants developed from 3/4PP endoderm with 2PAd mesenchyme showed similar PT formation efficiency to native associations (3/4PP and 3/4PA), suggesting a positive effect of 2PAd mesenchyme in PT formation (Table 1).

Our findings collectively demonstrate that local mesenchyme of the pharyngeal arches has the ability to regulate thymic potential in distinct pouches.

### PA mesenchymal signals that regulate PP thymic potential

To investigate the mesenchyme-derived signals that modulate thymus formation, we performed an unbiased, in-depth transcriptome analysis of isolated mesenchymal tissues from distinct PAs at cE2.5: 3/4PA, corresponding to a microenvironment favouring thymus formation; the dorsal region of 2PA (2Pad), which we demonstrated as having inhibitory properties to thymus development (Figure 5a). Total RNA was isolated from three biological replicate samples from the two mesenchyme regions and mRNA libraries were generated for paired-end illumina sequencing. High quality RNA-seq datasets were recovered for all samples, with an average of 7.7 million reads mapping uniquely to the *Gallus gallus* genome, corresponding to an average detection of ∼15 400 genes (Supplementary Dataset 1, figure 5b). Of these, 532 transcripts were found to be differentially expressed between 3/4PA and 2PAd with a log2FC >0.6 (adj pval <0.01) (complete list of genes in Supplementary Dataset 2).

**FIGURE 5.**
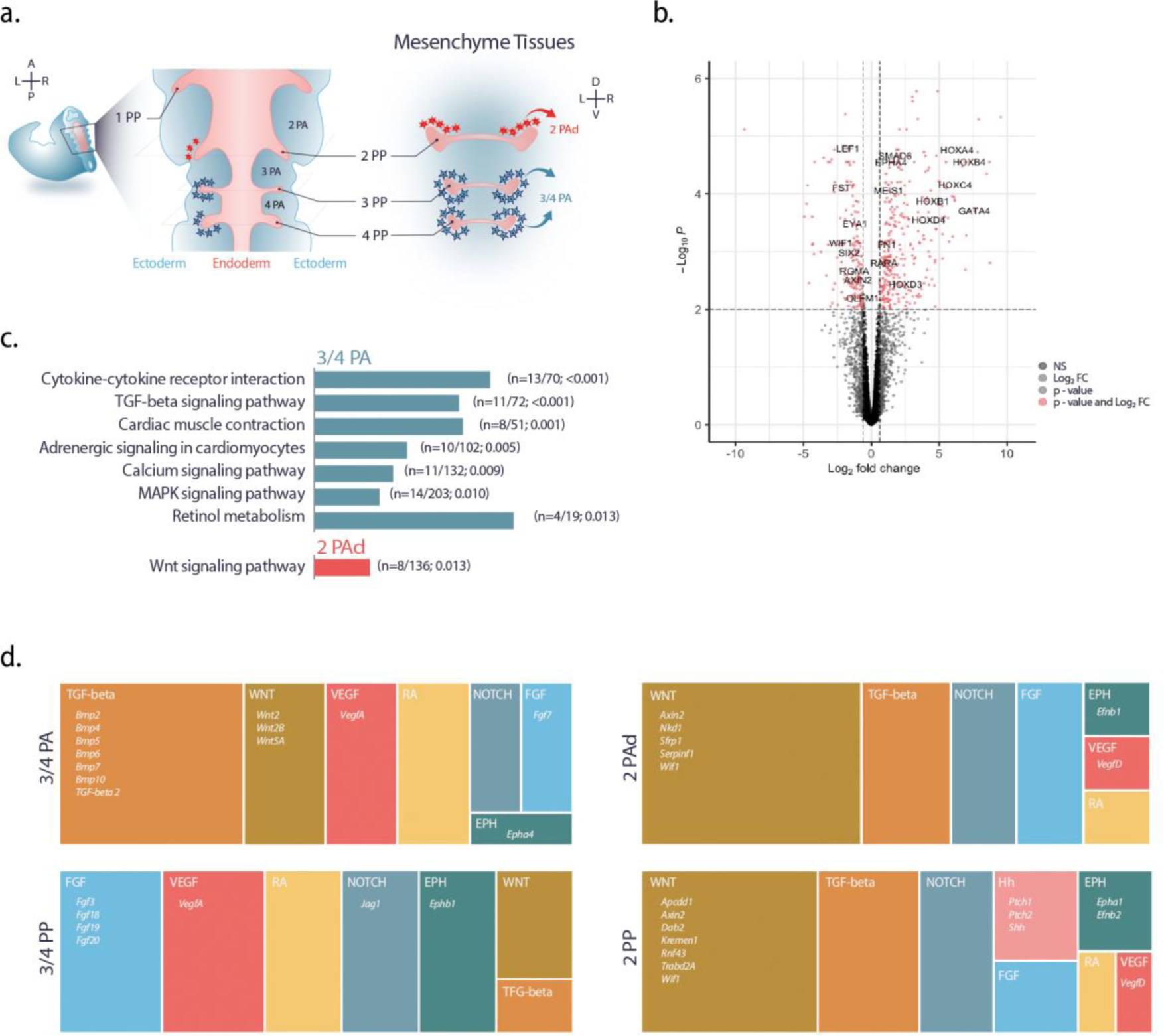
| Transcriptional profiles of pharyngeal mesenchyme tissues. a) Schematic representation of distinct mesenchyme tissues isolated from the embryonic pharyngeal region at cE2.5. b) Volcano plot showing the results of the differentially expressed analysis between 3/4PA and 2PAd mesenchyme. Red dots highlight transcripts passing the log2FC and adj pvalue cut-offs (adj pval <0.01 and |log2FC|>0.6); grey dots highlight transcripts passing only one of the cut-offs. d) Bar plot depicting KEGG pathway enrichment analysis for 532 genes differentially expressed in 3/4PA mesenchyme versus 2PAd mesenchyme (n= number of DE genes/total genes in pathway; adj pval). d) Tree-maps depict the number of up-regulated transcripts of major signalling pathways in 3/4PA mesenchyme, 2Pad mesenchyme, 3/4PP endoderm and 2PP endoderm transcriptomes. Inside the boxes a selection of genes relevant to details in the main text. A, Anterior; cE, chicken embryonic day; D, dorsal; EPH, EPH-Ephrin; FGF, fibroblast growth-factor; HH, Hedgehog; L, Left; P, Posterior; PA, Pharyngeal Arch; PAd, Dorsal region of Pharyngeal Arch; PP, Pharyngeal Pouch; R, right; RA, Retinoid Acid; V, Ventral; VEGF, Vascular endothelial growth-factor; TGF-beta, Transforming growth factor beta.

A closer inspection of functional gene annotation of DE genes revealed several transcripts encoding the Hox-family of genes, which may contribute to the A-P regionalization of PA mesenchyme. Although PA mesenchyme have multiple embryological sources and functions, the majority of cells derives from cardiac neural crest (CNC) [reviewed in (Dupin et al., 2006)]. CNC cells delaminate from distinct rhombomeres (R) with a specific Hox-code and migrate in a series of discrete streams to populate the pharyngeal arches [reviewed in (Parker and Krumlauf, 2020)]. In our analysis, we observed an upregulation of transcripts of genes encoding *Hoxa3-5*, *Hoxb1-4*, *Hoxc4*, and *Hoxd3-4* in the 3/4PA transcriptome (Figure 5b and Supplementary Dataset 2). Considering previous data [reviewed in (Parker and Krumlauf, 2020)], our observations suggest an unnoticed contribution of Hoxa5, Hoxb1, and Hoxc4 for arches posterior identity.

The study pursued with KEGG pathway enrichment analysis to improve our understanding to the function of upregulated transcripts detected in mesenchyme transcriptomes (Figure 5c and supplementary Table 1). We found an enrichment in Cytokine-cytokine receptor interaction and TGF-beta signalling pathway in 3/4PA mesenchymal transcriptome profile. Of note, several genes associated with these two signalling pathways were BMP ligands (supplementary Table 1) that have previously been identified as mesenchymal-derived signals involved in 3/4PP patterning and common T/PT primordium specification (Bleul and Boehm, 2005; Gordon et al., 2010; Jerome-majewska et al., 2002; Neves et al., 2012; Patel et al., 2006). In contrast, transcripts encoding genes associated to Wnt signalling pathway were only observed in 2PAd mesenchymal profile. Most of these genes (5 of 7) were pathway inhibitors, suggesting that they may operate as silencers for Wnt signalling at this embryonic stage in the dorsal region of the PA, similar to what is seen in later stages of thymus formation (Swann et al., 2017). This dataset analysis suggests specific modulation of signalling pathways according to PA mesenchymal settings, which may help to regulate the thymic potential of the endoderm of the pouches.

### Molecular crosstalk between PP endoderm and mesenchyme

To deepen our understanding of the signals involved in the distinct mesenchymal environments and respective epithelial-mesenchymal interactions, we performed manual examination of several signalling pathways components (EPH-Ephrin, FGF, Hedgehog, Notch, Retinoid Acid, TGF-beta, VEGF, and Wnt) using Go-term and Reactome-pathway enrichment (Supplementary Table 2). Treemaps generated with the number of DE genes between mesenchymal transcriptome profiles revealed distinct representativeness of the signalling pathways (Figure 5d). As in our previous KEGG pathway enrichment analysis (Figure 5c), herein the TGF-beta signalling was also the most represented pathway in 3/4PA mesenchyme. The second most representative set of up-regulated transcripts in the 3/4PA profile encoded Wnt signalling members (Supplementary Table 2). In fact, several of these were Wnt ligands, *Wnt2*, *Wnt2B*, and *Wnt5A*, suggesting that they could act as local mesenchymal-derived positive modulators of 3/4PP endoderm development and specification into TE cells. In a comparable developmental process, it was shown that Wnt5A secretion by dermal cells leads to autocrine *Fgf7* expression and consequent induction of *Foxn1* expression in hair follicular cells (Hu et al., 2010). Our data also revealed Fgf7 as the most enriched FGF-related transcript in 3/4PA mesenchyme, pointing to a similar Wnt5A/Fgf7/Foxn1 cascade in thymus induction at 3/4PP endoderm, probably mediated by fibroblast growth factor receptor R2-IIIb (Revest et al., 2001).

When examining the 2PAd mesenchyme’s upregulated transcripts, we noticed an inverse pattern in the predominance of signalling pathways. Similar to our previous KEGG pathway enrichment analysis (Figure 5c), Wnt signalling was the most representative pathway having enrichment of various upregulated transcripts encoding genes that serve as pathway inhibitors (Figure 5d and Supplementary Table 2).

To further assess the signals involved in epithelial-mesenchymal interactions that may govern thymus specification, we performed a similar methodological approach for the DE genes in PP endoderm transcriptomes (Figure 5d and Supplementary Table 3). Upregulated transcripts of 3/4PP showed a pattern of signalling pathways representation more similar to 3/4PA than 2PP, while the latter almost mirrored the 2PAd pattern. As in 2PAd profile, the Wnt signalling was the most represented pathway in 2PP endoderm transcriptome (Figure 5d) with several upregulated transcripts encoding Wnt inhibitors (Figure 5d and Supplementary Table 3). Interestingly, transcripts of a ligand and receptors of Hedgehog (Hh) signalling were enriched in the 2PP transcriptome but not in the 3/4PP (Figure 5d). The presence of active Hh signalling in 2PP endoderm was previously shown to repress Gcm2, restricting the development of PT to the caudal pouches (Grevellec et al., 2011). To avoid thymus development at the dorsal tip of the 2PP endoderm a similar scenario may be envisaged, along with a Wnt activity-restrictive environment in the dorsal region of the 2PA. In agreement, an ectopic expression of *Foxn1* was found in the dorsal tip of 2PP endoderm when Hh signalling was abolished (Figueiredo, 2011). The two most represented pathways in upregulated transcripts of 3/4PP endoderm were FGF (Figures 2c and 2d) and VEGF, followed *ex aequo* by EPH-Ephrin (EPH), Retinoid Acid (RA), and Notch signalling pathways (Figures 5d, 2c and 2d). A closer inspection of VEGF signalling showed an enrichment of *VegfA* transcripts in the epithelial-mesenchymal interactions 3/4PP-3/4PA, while *VegfD* transcripts were more expressed in the 2PP-2PAd environment. Similar to its functions in later stages of thymic development (Mü et al., 2005), VegfA may be required at this stage for vessel formation in the vicinity of the 3/4PP endoderm, a prerequisite for future colonization of the thymic rudiment by hematopoietic progenitor cells.

In the same way, we found enrichment of transcripts encoding unique ephrin receptors in the PP transcriptome profiles (Figure 5d and Supplementary Table 3), suggesting distinctive feature of Eph-ephrin interactions by potentially providing specific migratory routes while helping the establishment borders between different compartments in the arches. Ephrin signalling is known to participate in several thymus developmental scenarios [reviewed in (Muñoz et al., 2011)] and EfnB2 ligand was shown to be required for thymus migration during organogenesis (Foster et al., 2010).

The analysis of RA signalling in the 3/4PP transcriptomic profile revealed an enrichment of transcripts encoding for *RarB* and other pathway components (Figure 5d and Supplementary Table 3), along with the above-mentioned up-regulation of *Hox*-genes transcripts (Figures 3e and 3f). A similar gene expression landscape was observed for the 3/4PA transcriptomic profile (Figure 5b), suggesting that RA may acts as a caudalizing factor in a graded manner not only in pharyngeal endoderm (Bayha et al., 2009) but also in the surrounding arch mesenchyme. Given the observed upregulation of *Hoxb3 and Jag1* transcripts in 3/4PP transcriptomic we wonder whether Hoxb3-Jag1 signaling could be one of the defining events for 3/4PP endoderm identity and future specification in the TE. Notably, it was recently demonstrated that Notch ligand *Jag1* was specifically upregulated in the 2PA ectoderm when *Hoxb3* is ectopically expressed in Hoxb2 domain (Zhang et al., 2021).

As an attempt to further enhance our understanding of intercellular communication, we applied the unbiased computational bioinformatic NicheNet method, which predicts ligand– receptor pairs between interacting cells by combining their expression data with prior knowledge on signaling and gene regulatory networks (Browaeys et al., 2020). We used this methodology on the two 2PAd and 3/4PA mesenchymal compartments and infer active ligands and their gene regulatory effects on the two PP endoderm interacting cell populations, 2PP and 3/4PP, respectively. In our study, we opt to use only the upregulated genes in interacting PP endoderm receiver cells. The putative interactions that emerge from this strategy will portray a plausible scenario in which signalling pathways are implied to be activated by the expression of positive modulators and to be blocked by the expression of negative modulators, such as pathway inhibitors.

We started by prioritizing (with high regulatory potential score) the 20 ligands of each PA mesenchyme sender cells most likely to have affected the gene expression in interacting PP endoderm receiver cells (maps in Figure S3, schematic representation in Figure 6). When exploring 3/4PA-3/4PP interactions, we noticed that the GDF3 ligand showed a strong regulatory potential of several network genes in 3/4PP endoderm receiver cells (Figures 6 and S3a), while having the highest prior interaction potential with the receptor Tsukushi (Tsku) (Figures 6 and S3b). Tsku is a secreted protein that mediates multiple signalling pathways crucial for developmental processes such as cellular communication, proliferation, and cell fate determination [reviewed in (Istiaq and Ohta, 2022)]. However, the involvement of Tsku in thymus development remains unknown, as well as the putative Gdf3 (3/4PA) - Tsku (3/4PP) interaction, which may contribute to signalling during thymus specification and remains to be explored. In a complementary note to the above-mentioned, BMP ligands (Bmp4, Bmp7, and Bmp5) in 3/4PA mesenchyme showed a strong regulatory potential of the upregulated genes in 3/4PP endoderm (Figures 6 and S3a). The inferred ligand-target link was preferential with Bmpr2 in 3/4PP endoderm (Figures 6 and S3b), in accordance with the previously described expression of the receptor in this embryonic location (Neves et al., 2012). The observed chemokine-chemokine receptor interaction prediction, Cxcl12 (also named Sdf1) (3/4PA) - Cxcr4 (3/4PP), is concordant with the temporal-spatial role of Cxcr4-Cxcl12 signaling in developing pharynx. It was previously shown in chick and mouse embryos, that disruptions of Cxcr4 signaling in pharyngeal neural crest cells causes DiGeorge syndrome-like malformations including thymus hypoplasia (Escot et al., 2016).

**FIGURE 6.**
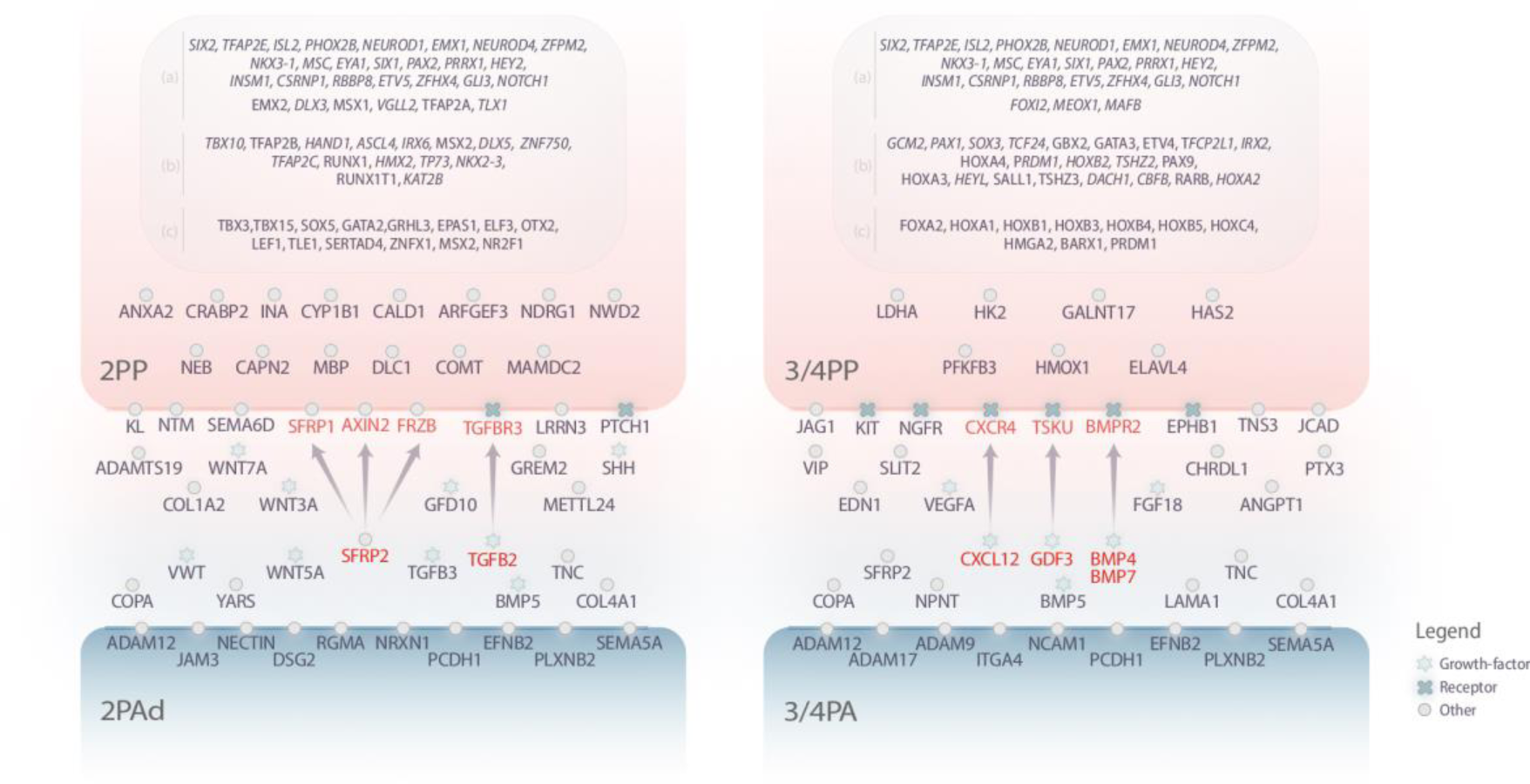
| Model of epithelial-mesenchymal interactions at early-stages of thymus development. Drawing representing cellular interactions between 2PPP endoderm – 2Pad mesenchyme and 3/4PP endoderm - 3/4PA mesenchyme. Molecular players involved in the crosstalk between PP endoderm and PA mesenchyme based on NicheNet analysis: 20 ligands (with high regulatory potential score) from each PA mesenchyme sender cells and upregulated gene networks in PP endoderm receiver cells. Arrows indicate ligand (in red) - target (in pink) interactions with the highest previous potential (details in the main text). TF&R expression landscape based on DE genes (in italic) analysis between endoderm tissues transcriptomes. a) Upregulated transcripts encoding TFs&Rs shared by 2PP and 3/4PP endoderm; b) Upregulated transcripts encoding TFs&Rs specific to each PP endoderm; c) Upregulated transcripts encoding TR&R belonging to the networks identified by NicheNet analysis.

When applying NicheNet analysis to 2PAd-2PP interactions, we observed a high regulatory potential score of Tgfb2 ligand to several network genes in 2PP endoderm receiver cells (Figures 6 and S3c), with a high score for prior interaction potential with three Tgfbrs (Figures 6 and S3d). It’s interesting to note that when comparing the putative interactions of Tgfb family members, the predicted Tgfb-Tgfbr interactions between 2PAd-2PP differ from those seen in the 3/4PA −3/4PP communication, where Bmp-Bmpr signaling is known to have a significant role. Another important difference in predicting active ligand–target links between 2PAd-2PP and 3/4PA −3/4PP conditions involved the ligand from Wnt signalling family, the secreted frizzled related protein 2 (Sfrp2). Although, Sfrp2 had the highest regulatory potential score in both conditions analysed, only in the 2PAd-2PP interactions Sfrp2 showed strong regulatory potential (Figures 6 and S3c) and prior interaction potential (Figures 6 and S3d) with other Wnt signalling inhibitors in 2PP endoderm (Axin2, Sfrp1, and Frzb). These interactions provide evidence in favour of the hypothesis that inhibition of Wnt signaling may help prevent thymus development at the dorsal tip of the 2PP endoderm.

Together, these findings provide some insight into possible pathways of cellular communication between 3/4PA-3/4PP and 2PAd-2PP that may help to regulate the thymic potential of PP endoderm.

## final remarks and conclusions

This study shows for the first time the capacity of non-canonical pouches endoderm to participate in thymus formation. We demonstrate the capacity of 2PP endoderm to differentiate into TE using the chick-quail chimera system. In human and mice, several hypotheses have been proposed to explain the existence of accessory cervical thymi, from migration defects of thymic rudiment to PT cellular transdifferentiation [reviewed in (Rodewald, 2008), (Li et al., 2013)]. Our data supports the hypothesis that a cervical thymus can arise independently of the thoracic thymus and from a distinct embryonic origin in the 2PP endoderm. In addition, we also observed ectopic PT formation with epithelium derived from the 2PP endoderm, extending the intrinsic potential for the development of a common T/PT primordium to non-canonical pouches.

Transcriptome profiling of the distinct endoderm territories in the pharyngeal arch region revealed a pool of shared upregulated TFs&Rs in the endoderm of canonical 3/4PP and non-canonical 2PP. Several of those were previously associated to thymus development (Bautista et al., 2021; Figueiredo et al., 2016; Li et al., 2003; Park et al., 2020), corroborating the existence of a common genetic program specifying thymic competence in endoderm progenitors of these pouches. A set of 17 TFs&Rs were for the first time associated to a likely conserved genetic program embodied in PP endoderm with thymic potential (Figure 6). Additionally, a specific Hox gene transcriptional profile was observed in the endoderm. The set of *Hox a1-a4*, *Hox b1-b5* and *Hoxc4* genes was identified as highly expressed in the endoderm of central pharynx and 3/4PP, suggesting its role in A-P regionalisation of the most posterior territories of developing pharynx. Of those, only *Hoxa2*, *Hoxa3* and *Hoxb2* transcripts were found to be DE in the 3/4PP endoderm in relation to the other two endoderm tissues. Given the functions and expression patterns previously described for the Hox gene family [reviewed in (Figueiredo et al., 2020), (Magaletta et al., 2022)], these genes may correspond to the minimal combination of Hox-genes mapping the posterior PP endoderm identity.

We further identified the mesenchymal cells that regulate PP endoderm thymic potential, showing that the 3/4PA mesenchyme promotes thymus formation whereas the dorsal region of the 2PA mesenchyme exerts an inhibitory effect. These distinct properties may be contextualised by considering the source of CNC delamination from the neural tube. In chicken development, 2PA is colonised by migratory CNC derived from rhombomere (R) 4 while the tip of 2PP is lined by CNC derived from R5 [reviewed by (Dupin et al., 2006)]. According to these findings, the dorsal and ventral domains of 2PP endoderm interact with different CNC sources, resulting in distinct epithelial-mesenchymal interactions at those pouch domains. A similar phenomenon is observed in 1PP, in which the dorsal portion is covered by cells migrating from R3 while remaining pouch territory is in contact with CNC migrating from R4. Interestingly, 3PA and 4PA are colonised by migratory CNC derived from R6-8, with no known specific final location within the arches [reviewed by (Dupin et al., 2006)].

The transcriptome profiles of 3/4PA and 2PAd mesenchyme were used to identify possible signals involved in the molecular crosstalk established with the PP endoderm, leading to the modulation of its potential. For this, we used available bioinformatic tools such as NicheNet that can predict which ligands influence the expression in another cell, which target genes are affected by each ligand and which signalling mediators may be involved (Browaeys et al., 2020). In this study we propose that a Wnt activity-restrictive environment along with activation of Hh signalling may be important to prevent thymus development at the dorsal tip of the 2PP endoderm (Figure 6). Additionally, the combined activity of Bmp, Fgf and RA signals seems to provide a proper environment for 3/4PP development, ultimately allowing thymus development (Figure 6). Experimental gain- and loss-of-functional strategies aiming at manipulating cell signalling in developing pharyngeal arches will provide the final confirmation of these concepts.

Overall, our data provide a better understanding of the molecular processes underlying interactions between pharyngeal pouch endoderm and pharyngeal arch mesenchyme that precede thymus formation. This information shall help not only to explore the signals and cellular properties that regulate thymus specification in the embryo, but also to design improved organoid systems that recapitulate epithelial-mesenchymal interactions and lead to efficient *ex vivo* generation of thymic organs for regenerative purposes.

## Materials and Methods

### Isolation of quail and chick embryonic tissues

Fertilised Japanese quail (*Coturnix coturnix japonica*) and chicken (*Gallus gallus*) eggs were incubated at 38 °C in a humidified incubator and embryos were dissected at specific times of development. Embryos were staged by microscopic examination according to Hamburger and Hamilton stages [HH, (Hamburger and Hamilton, 1992)] in the chick and to corresponding HH-stages in the quail.

Isolation of 2PP and 3/4PP endoderm was performed at embryonic day 3 (E3, HH-stage 21) of quail embryos and at E2.5 (HH-stage 19-20) of chick embryos, as previously described (Figueiredo and Neves, 2019; Le Douarin and Jotereau, 1975; Neves et al., 2012). Briefly, pharyngeal tissues were obtained by treating the wall of the embryonic pharynx with a solution of pancreatin (8 mg/ml, Sigma) for 30–90 min on ice, which allowed separation of pure endoderm from the pharyngeal mesenchyme. Mesenchymal tissues of E2.5–E3 (HH-stages 18–19) chick embryos were dissociated from endodermal and ectodermal tissues by enzymatic digestion with pancreatin using the same procedure described above. Somatopleura tissues were obtained from the embryonic territories at the level of somites 19–24 (Figueiredo and Neves, 2019).

The pharyngeal arch region (PAR) including 2PP/2PA to 4PP/4PA was dissected at qE3, as previously described (Figueiredo and Neves, 2018; Figueiredo and Neves, 2019).

### *In vitro* and *in ovo* tissue culture assays

The endoderm of 2PP and 3/4PP at qE3 were grown in association with mesenchymal tissues at cE2.5, as described (Figueiredo and Neves, 2019; Neves et al., 2012). Different sources of mesenchymal tissues were isolated: Somatopleura mesoderm (SM); 3/4PA mesenchyme; 2PA mesenchyme; ventral territory of 2PA mesenchyme (2PAv); and dorsal territory of 2PA mesenchyme (2PAd). In brief, 2–3 endodermal explants (2PP or 3/4PP endoderm) were combined with 2–3 mesenchymal explants on Nucleopore membrane filters (Millipore) supported by fine meshed metal grids (Goodfellows). The grids were then placed into culture dishes and partly immersed in RPMI-1640 (Sigma) supplemented with 10% FBS (Invitrogen) and 1X Pen/Strep (Invitrogen). The heterospecific associated tissues were cultured for 48 h at 37 °C in a humidified incubator containing 5% CO_2_. Following the incubation period, cultured tissues were either used for RNA isolation or grafted on the chorioallantoic membrane (CAM) of E8-chick embryos. The CAM behaves as a vascular supplier of nutrients and allows gas exchanges to grafted tissues, enabling their development *in ovo* for longer periods of time. Grafted tissues were allowed to further develop in ovo for 10 days in a humidified incubator at 38 °C, as described (Le Douarin and Jotereau, 1975; Neves et al., 2012).

PAR at qE3 were grown *in vitro* for 48h, as described (Figueiredo and Neves, 2018; Figueiredo and Neves, 2019). In brief, 6 PAR explants were placed on Nucleopore membrane filters (Millipore) supported by fine meshed metal grids (Goodfellows) and cultured as described above. Following the incubation period, cultured tissues were used for RNA isolation and qRT-PCR analysis.

### Quantitative real time RT-PCR

Total RNA from the samples was extracted using a combination of TRIzol reagent (Invitrogen) and RNeasy Mini Kit (QIAGEN) according to the manufacturer’s instructions. RNA samples were obtained from freshly isolated tissues at cE3: 2PP endoderm, 3/4PP endoderm and PAR; and, from *in vitro* cultured tissues for 48h: PAR and heterospecific associations of 3/4PP endoderm (qE3) with SM (cE2.5) or 2PAd mesenchyme (cE2.5) and heterospecific associations of 2PP endoderm (qE3) with SM (cE2.5). Each PP endoderm sample was collected from 15 to 20 embryos. Triplicates of each sample were generated for each condition. After DNAse treatment for 15 min, first-strand cDNA synthesis was performed in a total volume of 20μL, by reverse transcription of 300ng of total RNA using the SuperScript^TM^ III Reverse Transcriptase kit and Oligo (dT)_12-18_ Primer (Invitrogen), according to the manufacturer’s instructions. All steps of RNA extraction and cDNA synthesis were performed in a vertical laminar flow hood to avoid contamination. Concentration and purity of both the RNA and cDNA samples were determined using a NanoDrop® ND-1000 Spectrophotometer (Thermo Scientific).

To exclude the amplification of genomic DNA, primers were designed to span introns near the 3’poly-A tail using Primer3 software (Figueiredo et al., 2016). Quantitative RT-PCR (qRT-PCR) assays were run in a ViiA7^TM^ Fast Real-Time PCR System (Applied Biosystems) in MicroAmp® Optical 384-Well Reaction Plate (Applied Biosystems). Reactions were performed in a final volume of 10μL using 5μl of Power SYBR® Green PCR Master Mix (Applied Biosystems), 0.4μM final concentration of primers and 1μl (up to 1μg) of cDNA. Thermocycling conditions were as follows: an initial denaturation at 50°C for 20sec and 95°C for 10min, followed by 40 cycles at 95°C for 15sec and at 60°C for 1min. To confirm primer specificity, a melting curve was generated at the end of each experiment. Relative quantification of gene expression was determined by the ΔΔCt method (Livak and Schmittgen, 2001) using Hypoxanthine-guanine phosphoribosyltransferase (*Hprt*) as endogenous gene. Three technical replicates were used for each condition.

### Immunohistochemistry and *in situ* Hybridization

Chicken embryos and explants developed *in ovo* for 10 days were fixed overnight in 4% paraformaldehyde/PBS at 4 °C. Samples were then processed for immunohistochemistry and whole-mount *in situ* hybridisation.

Paraffin sections of CAM-explants were analysed by haematoxylin-eosin staining (H&E) to determine the number, size and morphology of thymic lobes and parathyroid glands formed. Sections of CAM-explants were further treated for immunocytochemistry with the QCPN antibody (for labelling of quail cells), CD3 antibody (Dako M725429-2, for labelling T-lymphoid cells) and anti-pan [Lu-5] Cytokeratin antibody (Pan CK) (Abcam; for labelling epithelial cells), as described (Figueiredo and Neves, 2018; Neves et al., 2012).

Whole-mount preparations were hybridised with *Fgf3*, *Fgf18,* and *Fgf19* probes as previously described (Etchevers et al., 2001; Henrique et al., 1995).

### Flow Cytometry

Thymi were isolated from chicken embryos at E17. cE17 roughly corresponds to the stage of development of the collected explants, that is qE13 (Ainsworth et al., 2010). Thymocytes suspensions were prepared by mechanically disrupting the thymus by gently pressing the tissue through a fine nylon mesh (pore size 70μm) to obtain a single cell suspension. The suspension was then purified by Ficoll density gradient separation (Ficoll-Paque PLUS GE Healthcare Life Sciences), to remove dead cells and red blood cells. All steps of the staining procedure were carried out on ice. Cells were stained using mouse anti-chicken mABs CD3-PE (clone CT-3; Southern Biotech) according to manufacturer’s instructions. Flow cytometry analysis was performed using LSR Fortessa cytometer (Becton Dickinson) and FlowJo software (TreeStar).

### Microscopy

H&E and immunohistochemistry images were collected using Software Leica Firewire and Leica DM2500 microscope with Leica DFC420 camera.

### RNA library preparation

Total RNA from the samples was extracted using a combination of TRIzol reagent (Invitrogen) and RNeasy Mini Kit (QIAGEN) according to the manufacturer’s instructions (Figueiredo et al., 2016). RNA samples were obtained from freshly isolated embryonic tissues of quail endoderm at E3 and chicken mesenchyme at E2.5. The isolated endoderm tissues were 2PP endoderm, 3/4PP endoderm and central pharynx region (Pharynx). The isolated mesenchyme tissues were 3/4PA mesenchyme and dorsal territory of 2PA mesenchyme (2PAd). Three replicates of 15-20 explants per sample were generated for each condition. Total RNA samples were validated for concentration and integrity (RIN ≥ 7) and 1 µg of total RNA was used to prepare libraries with the Stranded mRNA Library Preparation Kit according to manufacturer’s instructions. Libraries were sequenced on an Illumina Novaseq platform with paired end 150bp reads (acquired as a service to STAB Vida), with an average yield of 25M reads per sample (R1+R2). The raw RNA-Seq datasets are available through the European Nucleotide Archive under the study accession number PRJEB51508 (Gallus gallus dataset) and PRJEB51507 (C. coturnix dataset).

### RNA-Seq data analysis

Following quality assessment using FastQC version 0.11.5 (https://www.bioinformatics. babraham.ac.uk/projects/fastqc/), Cutadapt was used to remove sequencing adaptors and trim the first 10 nucleotides (Martin, 2011). The trimmed data was then filtered using in-house developed Perl script to remove reads with unknown nucleotides, homopolymers with length ≥50 nt or an average Phred score <30 (Amaral et al., 2014). Remaining reads, corresponding on average to 80% of the raw data, were aligned to the Ensemble genome references Coturnix_japonica_2.0 (GCA_001577835.1) or GRCg6a (GCA_000002315.5) for quail or chicken libraries, respectively. Alignment was performed using STAR version 2.5.0 with the following options: –outFilterType BySJout –alignSJoverhangMin 8 –alignSJDBoverhangMin 5 – alignIntronMax 100000 –outSAMtype BAM SortedByCoordinate –twopassMode Basic – outFilterScoreMinOverLread 0 –outFilterMatchNminOverLread 0 –outFilterMatchNmin 0 – outFilterMultimapNmax 1 –limitBAMsortRAM 10000000000 –quantMode GeneCounts (Dobin et al., 2013). Gene counts were determined using the htseq-count function from HTseq (version 0.9.1) (Anders et al., 2015) in union mode and discarding low quality score alignments (–a 10), using the Ensembl Coturnix_japonica_2.0 or GRCg6a genome annotations. On average, 67% and 77% of the filtered reads from quail and chicken samples mapped to a single genomic location, corresponding to ∼14 500 and ∼15 400 detected genes, respectively. Supplementary Dataset 1 presents the summary information of the RNA-seq datasets.

Clustering of normalized gene counts and Principal Component Analysis (PCA) for exploratory data analysis were performed with the pheatmap R package using the euclidean distance matrix computation of the dist function, and the ggfortify:plotPCA function, respectively.

Differential Expression Analysis (DEA) for RNA-Seq gene counts was performed with the limma Bioconductor package using the voom method to convert the read-counts to log2-cpm, with associated weights, for linear modeling (Law et al., 2014; Ritchie et al., 2015). DEA was performed by making all possible comparisons between the three C. japonica experimental conditions or by comparing the two G. gallus experimental conditions, using all available replicate data. Genes showing up or down-regulation with an adjusted *p value* <0.01 and a log2 fold change (log2FC) of |log2FC|>0.6 were considered differentially expressed (Supplementary Dataset 2).

Heatmaps for DE genes were generated using the pheatmap function from the R pheatmap package with the ward.D clustering method. Volcano plots were produced with the EnchancedVolcano R package.

### Functional enrichment and ligand-receptor interaction analysis

The Biomart annotation for Coturnix japonica and the Manteia database Public release v. 7.0 October 2018 annotation for Gallus gallus (http://manteia.igbmc.fr/index.php) were used for ENSEMBL to Gene Symbol identifier conversions, with manual correction for ambiguous or missing annotations of Hox and ligand/receptor genes.

Functional enrichment analysis was performed with the GOFuncR R package. Heatmap representation of selected enriched GO terms was generated with the pheatmap R package. Cluster identification was performed using the pheatmap package cutree function with k=6.

The DE genes in PA mesenchymal transcriptomes were mapped to the KEGG database and searched for significantly enriched KEGG pathways at *p* < .05 level.

Receptor ligand-interaction networks were identified using the NicheNet R package and corresponding ligand-target matrix and ligand-receptor and weighted network datasets (Browaeys et al., 2020). 3/4PA cells were defined as senders, 3/4PP cells as receivers, and upregulated genes in 3/4PP versus 2PP as genes of interest. Alternatively, 2PAd cells were considered as senders, 2PP cells as receivers, and upregulated genes in 2PP versus 3/4PP as genes of interest. Gene expression datasets were filtered for a minimum log expression level >4, and analysis focused on the top 20 prioritized ligands. All other parameters were used as default.

### Statistical analyses

Analyses were performed using statistical software (GraphPad Prism 7.00; GraphPad Software). Two-tailed Student’s *t*-tests were used for qRT-PCR analysis. Results were considered significantly different when the *P* value was less than 0.05 (*P < 0.05, **P < 0.001, ***P < 0.0001).

## Acknowledgments

The authors are grateful to Leonor Parreira and Georg Hollander for helping raise this project from the ground and to Rita Zilhão and António Cidadão for useful discussions. We are also thankful to Solveig Thorsteinsdottir and Edgar Gomes for reagents and Interaves Portugal for contributing with quail fertilised eggs. This work was supported by Fundação para a Ciência e para a Tecnologia (FCT), project PTDC/SAU-BID/115264/2009 and Leonor Magalhães received two research grants from Gabinete de Apoio à Investigação Científica, Tecnológica e Inovação (GAPIC) – Faculdade de Medicina da Universidade de Lisboa. Work in MGC’s lab is supported by FCT, Portugal through grants UIDB/04046/2020 and UIDP/04046/2020 Centre grants from FCT (to BioISI).

## DATA AVAILABILITY

The 30 raw fastq files of the RNA-seq data generated for this study have been submitted to the European Nucleotide Archive under the umbrella project “Thymus inception” (PRJEB51509), with individual study accession numbers PRJEB51507 (quail dataset) and PRJEB51508 (chicken dataset). The microarray data set is hosted in the open data repository Zenodo (doi 10.5281/zenodo.8155337).

## AUTHOR CONTRIBUTIONS

HN contributed to the conceptualization, design of experiments, experiment execution, analysis of results and scientific illustrations. IA and LM carried out the experiments. MGC contributed to the experimental design of the transcriptome profiling, performed the RNA-seq data analysis and analysis of results. VP contributed with technical support. DH contributed to the analysis of results. HN and DH wrote the manuscript.

