## Supplementary Figures and Tables for "Thymus formation in uncharted embryonic territories"

a.

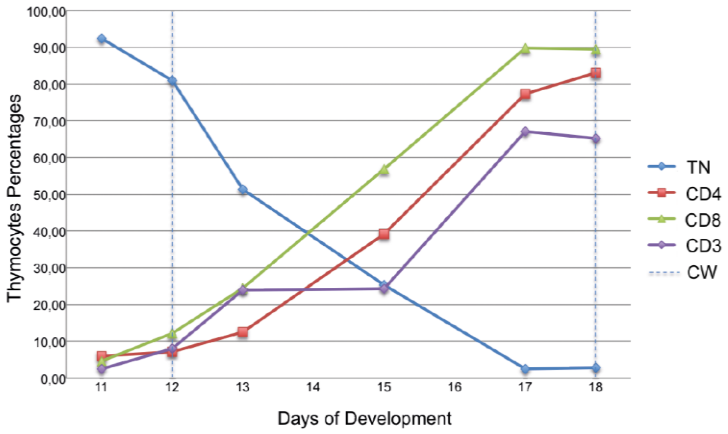

100,00
90,00
80,00
70,00
60,00
50,00
40,00
30,00
20,00
10,00
0,00
Thymocytes Percentages
11
12
13
14
15
16
17
18
Days of Development
TN
CD4
CD8
CD3
CW


Figure S1.

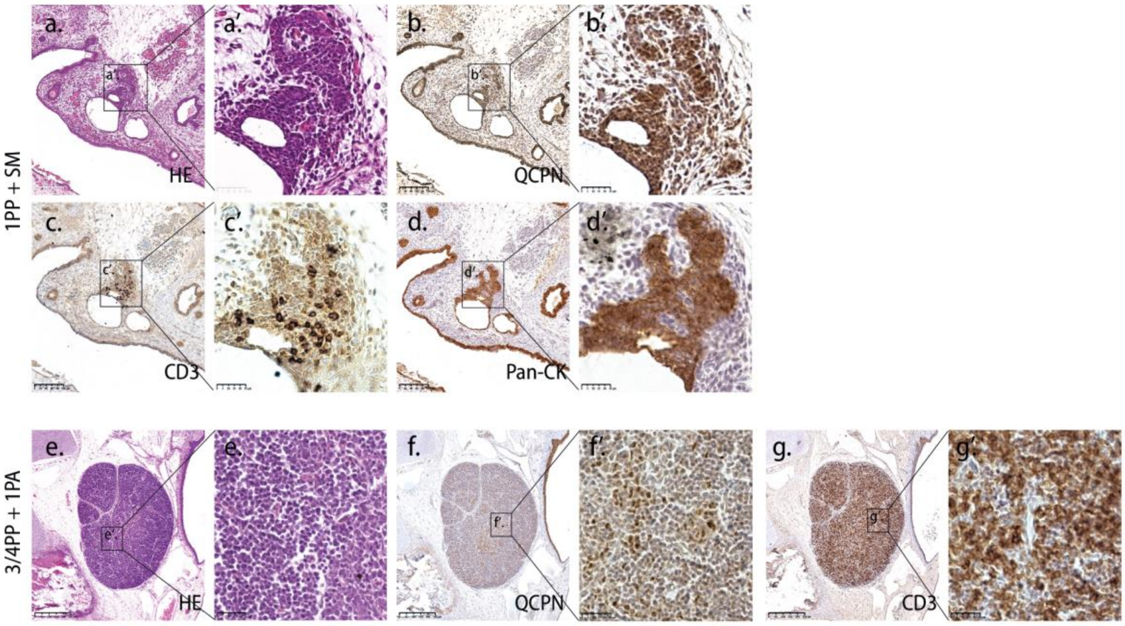

a.
a'.
a'
HE
b.
b'.
b'
QCPN
c.
c'.
c'
CD3
d.
d'.
d'
Pan-CK
1PP + SM
e.
e'.
e'
HE
f.
f'.
f'
QCPN
g.
g'.
g'
CD3
3/4PP + 1PA


Figure S2.

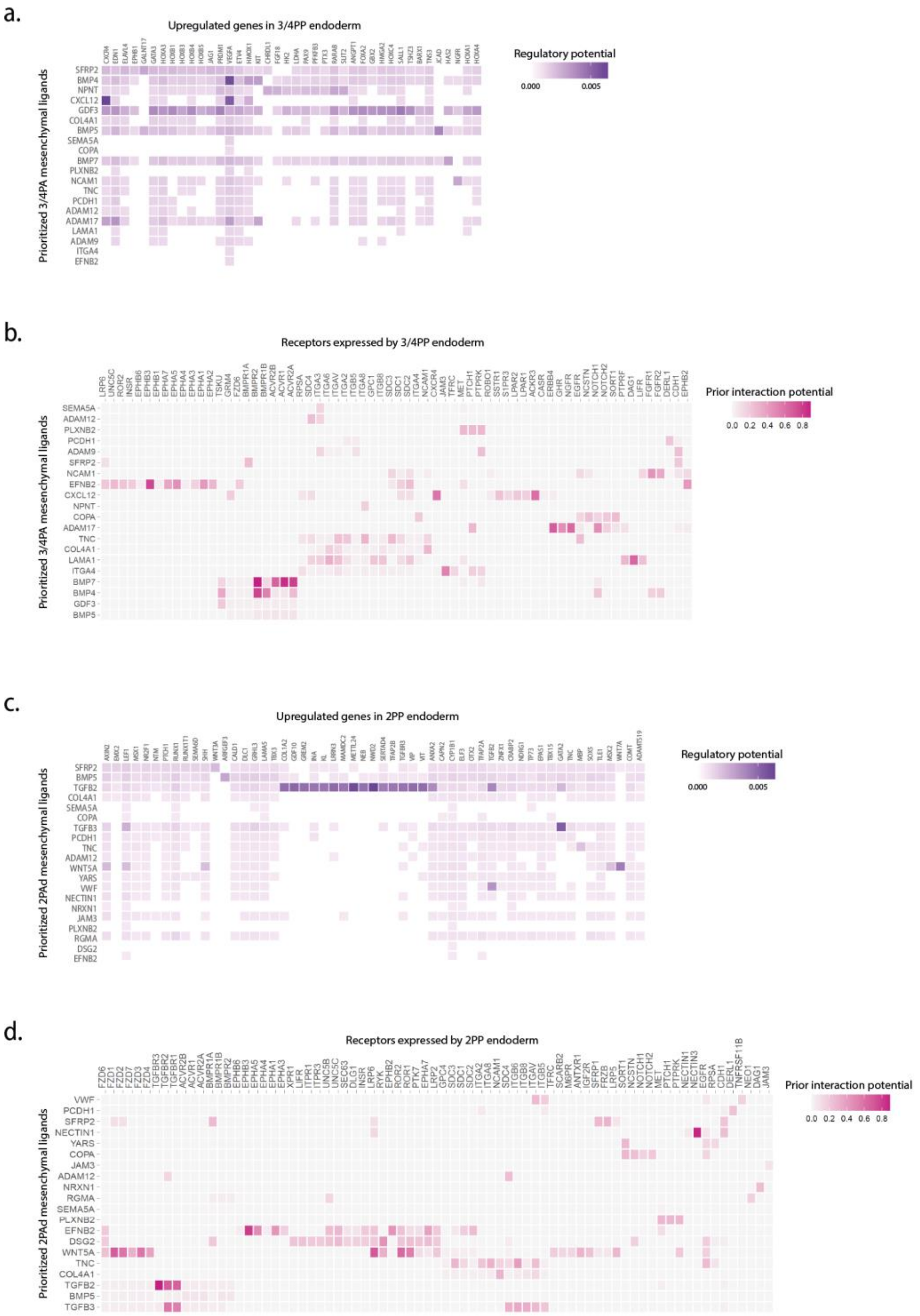
a.
Upregulated genes in 3/4PP endoderm
Prioritized 3/4PA mesenchymal ligands
Regulatory potential
0.000
0.005
SFRP2
BMP4
NPNT
CXCL12
GDF3
COL4A1
BMP5
SEMA5A
COPA
BMP7
PLXNB2
NCAM1
TNC
PCDH1
ADAM12
ADAM17
LAMA1
ADAM9
ITGA4
EFNB2
b.
Receptors expressed by 3/4PP endoderm
Prioritized 3/4PA mesenchymal ligands
Prior interaction potential
0.0 0.2 0.4 0.6 0.8
SEMA5A
ADAM12
PLXNB2
PCDH1
ADAM9
SFRP2
NCAM1
EFNB2
CXCL12
NPNT
COPA
ADAM17
TNC
COL4A1
LAMA1
ITGA4
BMP7
BMP4
GDF3
BMP5
c.
Upregulated genes in 2PP endoderm
Prioritized 2PAd mesenchymal ligands
Regulatory potential
0.000
0.005
SFRP2
BMP5
TGFB2
COL4A1
SEMA5A
COPA
TGFB3
PCDH1
TNC
ADAM12
WNT5A
YARS
VWF
NECTIN1
NRXN1
JAM3
PLXNB2
RGMA
DSG2
EFNB2
d.
Receptors expressed by 2PP endoderm
Prioritized 2PAd mesenchymal ligands
Prior interaction potential
0.0 0.2 0.4 0.6 0.8
VWF
PCDH1
SFRP2
NECTIN1
YARS
COPA
JAM3
ADAM12
NRXN1
RGMA
SEMA5A
PLXNB2
EFNB2
DSG2
WNT5A
TNC
COL4A1
TGFB2
BMP5
TGFB3

Figure S3.

### LEGENDS TO SUPPLEMENTARY FIGURES

**Supplementary Figure S1| T-cell differentiation during chicken embryonic development.** Flow cytometry analysis of thymocytes harvested from E11, E12, E13, E15, E17 and E18 thymi. Cells were stained using a panel of mouse anti-chicken monoclonal antibodies from Southern Biotech: anti-CD4 FITC (clone CT-4), anti-CD8 FITC (clone CT-8), anti-CD8 PE (clone CT-8) and anti-CD3 PE (clone CT-3) according to manufacturer's instructions. TN, Triple Negative thymocytes (CD3-CD4-CD8-); CW, Colonization Waves by LPCs (2nd and 3rd CW at E12 and E18, respectively); E – embryonic day.

**Supplementary Figure S2 | Thymus formation in CAM-derived explants of PP endoderm and mesenchyme associations.** Serial sections of thymi stained with H&E (a and e), and immunodetected with QCPN (quail cells) (b and f), CD3 (T-cells) (c and g) and anti-pan cytokeratin (CK) antibody (epithelial cell marker) (d) antibodies. Top panel (a-d) - thymus derived from associations of quail 1PP endoderm with chicken SM. Bottom panel (e-g) - A thymus derived from associations of quail 3/4PP endoderm with chicken 1PA mesenchyme. Arrowheads point immunostaining positive cells for QCPN (quail thymic epithelium) and for CD3 (chicken T-cells differentiated in the thymus). CAM, chorioallantoic membrane; E, embryonic day; PA, Pharyngeal Arch; PP, Pharyngeal Pouch; SM, somatopleura mesoderm. Scale bars, 100μm.

**Supplementary Figure S3 |NicheNet analyses of ligand-receptor interactions between PA mesenchyme -PP endoderm.** Ligand-target matrix denoting the regulatory potential between the 3/4PA mesenchymal cells (senders) and upregulated genes in endoderm of 3/4PP (receivers) (a). Ligand-target matrix denoting prior interaction potential between the 3/4PA mesenchymal cells (senders) and receptors in endoderm of 3/4PP (receivers)(b). Ligand-target matrix denoting the regulatory potential between the 2PAd mesenchymal cells (senders) and upregulated genes in endoderm of 2PP (receivers)(c). Ligand-target matrix denoting prior interaction potential between the 2PAd mesenchymal cells (senders) and receptors in endoderm of 2PP (receivers)(d). Gene expression datasets were filtered for a minimum log expression level >4, and analysis focused on the top 20 prioritized ligands.

### Supplementary Tables

Supplementary Table 1. **KEGG pathway analysis of PA mesenchymal transcriptomes**

| | Pathway | Genes |
|---|---|---|
| **3/4PA mesenchyme** | **Cytokine-cytokine receptor interaction** | BMP10, cVg1, FAS, BMP7, TGFB2, BMP4, BMP6, BMP5, BMP2, INHBA, CCR7, GDF6, TNFRSF11B |
| | **TGF-beta signaling pathway** | BMP7, TGFB2, BMP4, BMP6, BMP5, SMAD6, BMP2, INHBA, GDF6, BAMBI, PITX2 |
| | **Cardiac muscle contraction** | TNNT2, TNNC1, TPM4, MYL2, CACNA1D, MYL3, ACTC1, RYR2 |
| | **Adrenergic signaling in cardiomyocytes** | TNNT2, TNNC1, TPM4, MYL2, CACNA1D, MYL3, ACTC1, RYR2, MAPK12, MAPK11 |
| | **Calcium signaling pathway** | TNNC1, PDGF-A, MYLK3, CACNA1D, PDGFC, VEGFA, SLC25A4, RYR2, KDR, FGFR3, Fgf7 |
| | **MAPK signaling pathway** | PAK1, HSPB1, PDGF-A, CACNA1D, FAS, PDGFC, TGFB2, VEGFA, MKP3, KDR, FGFR3, MAPK12, Fgf7, MAPK11 |
| | **Retinol metabolism** | ALDH1A2, DHRS3, CYP26A1, RDH10 |
| **2PAd mesenchyme** | **Wnt signaling pathway** | SERPINF1, SFRP1, NKD1, TLE1, LEF1, AXIN2, WIF1, LGR4 |

Supplementary Table 2. **Genes of main signalling pathways differentially expressed in PA mesenchyme.**

| | 3/4PA mesenchyme | | 2PAd mesenchyme | |
|---|---|---|---|---|
| **VEGF signaling** (R-HSA-194138) | CYFIP2 | Cytoplasmic FMR1-interacting protein 2 | VEGFD | Vascular endothelial growth factor D |
| | HSPB1 | Heat shock protein beta-1 | | |
| | MAPK11 | Mitogen-activated protein kinase 11 | | |
| | MAPK12 | Mitogen-activated protein kinase 12 | | |
| | PAK1 | Serine/threonine-protein kinase PAK 1 | | |
| | KDR | Vascular endothelial growth factor receptor | | |
| | VEGFA | Vascular endothelial growth factor A | | |
| **Retinoic Ac. signaling** (R-HSA-5362517) | ALDH1A2 | Retinal dehydrogenase 2 | CYP26C1 | Cytochrome P450 26C1 |
| | CYP26A1 | Cytochrome P450 26A1 | | |
| | CRABP1 | Cellular retinoic acid-binding protein 1 | | |
| | DHRS3 | Short-chain dehydrogenase/reductase 3 | | |
| | RARA | Retinoic acid receptor alpha | | |
| | RBP5 | Retinol Binding Protein 5 | | |
| | RDH10 | Retinol dehydrogenase 10 | | |
| **TGFB / BMP signaling** (R-HSA-9006936; GO:0030509) | BAMBI | BMP and activin membrane-bound inhibitor homolog | CHRDL1 | Chordin-like protein 1 |
| | BMP2,4,5,6,7,10 | Bone morphogenetic protein 2, 4, 5, 6, 7 and 10 | COL1A2 | Collagen alpha-2 (I) chain |
| | CGN | Cingulin | FST | Follistatin |
| | DLX5 | Homeobox protein DLX-5 | TGFBR2 | TGF-beta receptor type-2 |
| | FKBP1B | Peptidyl-prolyl cis-trans isomerase FKBP1A | | |
| | GDF6 | Growth/differentiation factor 6 | | |
| | GPC3 | Glypican-3 | | |
| | INHBA | Inhibin beta A chain | | |
| | PITX2 | Paired like homeodomain 2 | | |
| | Q9W6I1 | RNA-binding protein involved in the regulation of smooth muscle cell differentiation and proliferation in the | | |
| | SMAD6 | Mothers against decapentaplegic homolog 6 | | |
| | SULF1 | Sulfatase 1 | | |
| | TGFB2 | Transforming growth factor beta-2 | | |
| **Wnt signaling** (R-HSA-195721; GO:0016055) | CTNND2 | Catenin delta-2 | AXIN2 | Axin-2 |
| | DKK3 | Dickkopf-related protein 3 | DIXDC1 | Dixin |
| | FZD8 | Frizzled-8 | LEF1 | Lymphoid enhancer-binding factor 1 |
| | PRICKLE1 | Prickle-like protein 1 | LGR4 | Leucine-rich repeat-containing G-protein coupled |
| | WNT2, 2B, 5A | Protein Wnt-2, 2b and 5a | NKD1 | Naked cuticle homolog 1 |
| | SFRP2 | Secreted frizzled-related protein 2 | WIF1 | Wnt inhibitory factor 1 |
| | | | SERPINF1 | Serpin family F member 1 |
| | | | SFRP1 | Secreted frizzled-related protein 1 |
| | | | SOX9 | Transcription factor SOX-9 |
| | | | TLE1 | Transducin-like enhancer protein 1 |
| **Notch signaling** ( R-HSA-157118; GO:0007219) | HDAC10 | Histone Deacetylase 10 | HDAC9 | Histone deacetylase 9 |
| | HES5 | Transcription factor HES-5 | SNAI2 | Zinc finger protein SNAI2 |
| | HOXD3 | Homeobox protein Hox-D3 | TLE1 | Transducin-like enhancer protein 1 |
| | NEURL1B | Neuralized E3 Ubiquitin Protein Ligase 1B | | |
| **EPH-Ephrin signaling** (R-HSA-2682334) | EPHA4 | Ephrin type-A receptor 4 | EFNB1 | Ephrin-B1 |
| | PAK1 | Serine/threonine-protein kinase PAK 1 | | |
| **FGF signaling** (R-HSA-190236) | FGF7 | Fibroblast growth factor 7 | FGF19 | Fibroblast growth factor 19 |
| | FGFRL1 | Fibroblast growth factor receptor-like 1 | FGFR3 | Fibroblast growth factor receptor 3 |
| | FGFR3 | Fibroblast growth factor receptor 3 | FLRT3 | fibronectin leucine rich transmembrane protein 3 |
| | SPRY2 | Protein sprouty homolog 2 | | |

List of genes depicted by alphabetic order.

Supplementary Table 3. **Genes of main signalling pathways differentially expressed in PP endoderm.**

| | 3/4PP endoderm | | 2PP endoderm | |
|---|---|---|---|---|
| **VEGF signaling** (R-HSA-194138) | NRP1 | Neuropilin-1 | PIK3R1 | Phosphatidylinositol 3-kinase regulatory subunit alpha |
| | PIK3CB | Phosphatidylinositol 4,5-bisphosphate 3-kinase catalytic subunit beta isoform | VEGFD | Vascular endothelial growth factor D |
| | PTK2 | Focal adhesion kinase 2 | | |
| | VEGFA | Vascular endothelial growth factor A | | |
| **Retinoic Ac. signaling** (R-HSA-5362517) | CRABP1 | Cellular retinoic acid-binding protein 1 | CYP26C1 | Cytochrome P450 26C1 |
| | RARB | Retinoic acid receptor beta | CRABP2 | Cellular retinoic acid binding protein 2 |
| | RDH10 | Retinol dehydrogenase 10 | | |
| **TGFB/BMP signaling** (R-HSA-9006936; GO:0030509) | CHRDL1 | Chordin-like protein 1 | BAMBI | BMP and activin membrane-bound inhibitor homolog |
| | | | COL1A2 | Collagen alpha-2(I) chain |
| | | | FST | Follistatin |
| | | | GDF6 | Growth/differentiation factor 6 |
| | | | GREM2 | Gremlin-2 |
| | | | ITGB6 | Integrin beta-6 |
| | | | SOSTDC1 | Sclerostin domain-containing protein 1 |
| | | | TGFB2, 3 | Transforming growth factor beta-2 and -3 proproteins |
| | | | TGFBR3 | Transforming growth factor beta receptor type 3 |
| | | | WWTR1 | WW domain-containing transcription regulator protein 1 |
| **Wnt signaling** (R-HSA-195721; GO:0016055) | LGR5 | Leucine-rich repeat-containing G-protein coupled receptor 5 | APCDD1 | APC down-regulated 1 |
| | SOX3 | Transcription factor SOX-3 | AXIN2 | Axin-2 |
| | | | DAB2 | Disabled homolog 2 |
| | | | FZD10 | Frizzled-10 |
| | | | KREMEN1 | kringle containing transmembrane protein 1 |
| | | | LEF1 | Lymphoid enhancer-binding factor 1 |
| | | | NXN | Nucleoredoxin |
| | | | RNF43 | Ring finger protein 43 |
| | | | ROR1 | Inactive tyrosine-protein kinase transmembrane receptor ROR1 |
| | | | WIF1 | Wnt inhibitory factor 1 |
| | | | WNT2B, 3A, 6, 7A, 7B, 9B | Protein Wnt-2b, 3a, 6, 7a, 7b and 9b |
| | | | TLE1 | Transducin-like enhancer protein 1 |
| | | | TRABD2A | TraB domain containing 2A |
| | | | SOX9 | Transcription factor SOX-9 |
| **Notch signaling** ( R-HSA-157118; GO:0007219) | FABP7 | Fatty acid binding protein 7 | ELF3 | ETS-related transcription factor Elf-3 |
| | HEYL | Hairy/enhancer-of-split related with YRPW motif-like protein | FBXW7 | F-box/WD repeat-containing protein 7 |
| | JAG1 | Protein jagged-1 | HDAC9 | Histone deacetylase 9 |
| | | | KAT2B | Histone acetyltransferase KAT2B |
| | | | RUNX1 | Runt-related transcription factor 1 |
| | | | SNAI2 | Zinc finger protein SNAI2 |
| | | | TLE1 | Transducin-like enhancer protein 1 |
| | | | TP63 | Tumor protein 63 |
| **EPH-Ephrin signaling** (R-HSA-2682334) | DNM1 | Dynamin-1 | EFNB2 | Ephrin-B2 |
| | EPHB1 | Ephrin type-B receptor 1 | EPHA1 | Ephrin type-A receptor 1 |
| | PTK2 | Focal adhesion kinase 2 | KALRN | kalirin RhoGEF kinase |
| | | | NGEF | neuronal guanine nucleotide exchange factor |
| **FGF signaling** (R-HSA-190236) | FGF3,18, 19, 20 | Fibroblast growth factor 3, 18, 19 and 20 | ANOS1 | Anosmin-1 |
| | | | FGFR2 | Fibroblast growth factor receptor 2 |
| | | | KL | Klotho |
| | | | PIK3R1 | Phosphatidylinositol 3-kinase regulatory subunit alpha |
| **Hedgehog signaling** (R-HSA-5358351) | | | ADCY5 | Adenylate cyclase type 5 |
| | | | HHIP | Hedgehog-interacting protein |
| | | | PTCH1 | Protein patched homolog 1 |
| | | | PTCH2 | Protein patched homolog 2 |
| | | | SHH | Sonic hedgehog protein |

List of genes depicted by alphabetic order.
